# NOD2 restrains probiotic-mediated TLR2-dependent antitumor immunity

**DOI:** 10.64898/2026.10.08.757712

**Authors:** Xing Zhang, Natavan Dudkina, Kyong T. Fam, Stanislav Dikiy, Adrianna M. Turner, Anh Vu-Nguyen, Bao Anh Nguyen, Tingting Dai, Howard C. Hang

## Abstract

The gut microbiota influences cancer immunotherapy, but how interactions among host microbial sensors determine the activity of individual bacteria remains unclear. Here, we show that NOD2 restrains TLR2-dependent antitumor immunity induced by *Lactiplantibacillus plantarum*. Oral administration of live or heat-killed *L. plantarum* enhanced immune checkpoint blockade efficacy in mouse tumor models. This activity required host TLR2 signaling and was diminished by disruption of bacterial lipoprotein. In contrast, NOD2 deficiency enhanced tumor control, intratumoral interferon-associated programs, and T cell activation. NOD2 deficiency also amplified chemokine and interferon-associated responses to *L. plantarum* and the synthetic TLR2 agonist Pam3CSK4 in myeloid cells. Genetic loss or pharmacological inhibition of NOD2 improved Pam3CSK4-mediated tumor control in combination with checkpoint blockade. These findings identify NOD2 as a functional checkpoint on TLR2-driven antitumor immunity and demonstrate that interactions between host microbial-sensing pathways shape the therapeutic activity of commensal bacteria and innate immune agonists.

**Highlights:**

- *L. plantarum* enhances checkpoint blockade through host TLR2
- Bacterial lipoproteins contribute to antitumor activity
- NOD2 restrains inflammatory and antitumor responses to *L. plantarum*
- Genetic loss or pharmacological inhibition of NOD2 improves TLR2 agonist efficacy

## Introduction

Responses to immune checkpoint blockade (ICB) vary considerably among patients, and many do not achieve durable clinical benefit. The gut microbiota has emerged as an important determinant of ICB efficacy, with multiple bacterial species enriched in responding patients and several shown to enhance antitumor immunity in preclinical models^1–5^. However, the molecular interactions linking individual commensal bacteria to host immune pathways remain incompletely defined. In addition, individual microbes carry multiple immunostimulatory molecules that can engage different host receptors simultaneously. How these signals interact across innate immune pathways to shape the antitumor activity of a given commensal remains poorly understood.

Host genetic variation provides an additional layer of complexity to antitumor immunity and ICB response. Germline variation in HLA class I genes has been associated with clinical outcomes following checkpoint blockade^6^, while inherited variation in immune pathways, including IFIH1, STING1, and PRKCD, has been linked to tumor immune states and antitumor immunity^7,8^. Notably, loss-of-function variants in NOD2, among the strongest genetic risk factors for Crohn’s disease^9,10^, have also been associated with improved ICB outcomes in patients with non-small cell lung cancer (NSCLC)^11^. However, this association was not reproduced in an independent cohort^12^, suggesting that the impact of NOD2 genotype on antitumor immunity may be context dependent. These studies have largely considered host genotype independently of the microbiota. How host genetic variations in microbial sensing pathways interact with specific commensal bacteria to shape antitumor immunity and ICB efficacy therefore remains unresolved.

Here, we investigate how host genetic variation in microbial sensing shapes the antitumor activity of a commensal bacterium. *Lactobacillus* species are Gram-positive lactic acid bacteria that are found in animals, plants, and human microbiota as well as in fermented foods^13^ with relevance in agriculture, human health, and medicine. As probiotics^14^, *Lactobacillus* species have diverse metabolic activity and are implicated in colonization resistance against pathogens and enhanced intestinal barrier function. Using *Lactiplantibacillus plantarum ATCC 8014* as a model commensal, we identified opposing roles for TLR2 and NOD2 in bacterial-mediated tumor control. *L. plantarum* antitumor activity required host TLR2 and was diminished by disruption of bacterial lipoprotein, whereas disruption of capsular polysaccharide biosynthesis had little effect. In contrast, NOD2 deficiency enhanced tumor control and amplified chemokine and interferon-associated responses to *L. plantarum* and the TLR2 agonist Pam3CSK4. Genetic loss or pharmacological inhibition of NOD2 also enhanced TLR2 agonist-mediated antitumor activity. Together, these findings identify bacterial lipoprotein–TLR2 signaling as a mechanism contributing to *L. plantarum*-mediated cancer immunotherapy and reveal NOD2 as a host checkpoint that restrains this response.

## Results

### *L. plantarum* potentiates immune checkpoint blockade

We previously showed that *Enterococcus faecium* enhances ICB through the peptidoglycan hydrolase SagA^15,16^. To determine whether SagA-associated immunostimulatory activity extends beyond *Enterococcus*, we examined a representative panel of phylogenetically and ecologically diverse lactobacilli harboring SagA orthologs, including species frequently detected in the human gastrointestinal tract and/or widely used as probiotics. Culture supernatants from all species tested activated NOD2 in HEK-Blue-based NOD2 reporter cells **(Fig. S1A)**. However, among live bacteria, only *Lactiplantibacillus plantarum* induced detectable NOD2 activation **(Fig. 1A)**, whereas none activated NOD1 **(Fig. S1B)**. These findings prompted us to investigate the immunomodulatory activity of *L. plantarum*.

**Figure 1.**
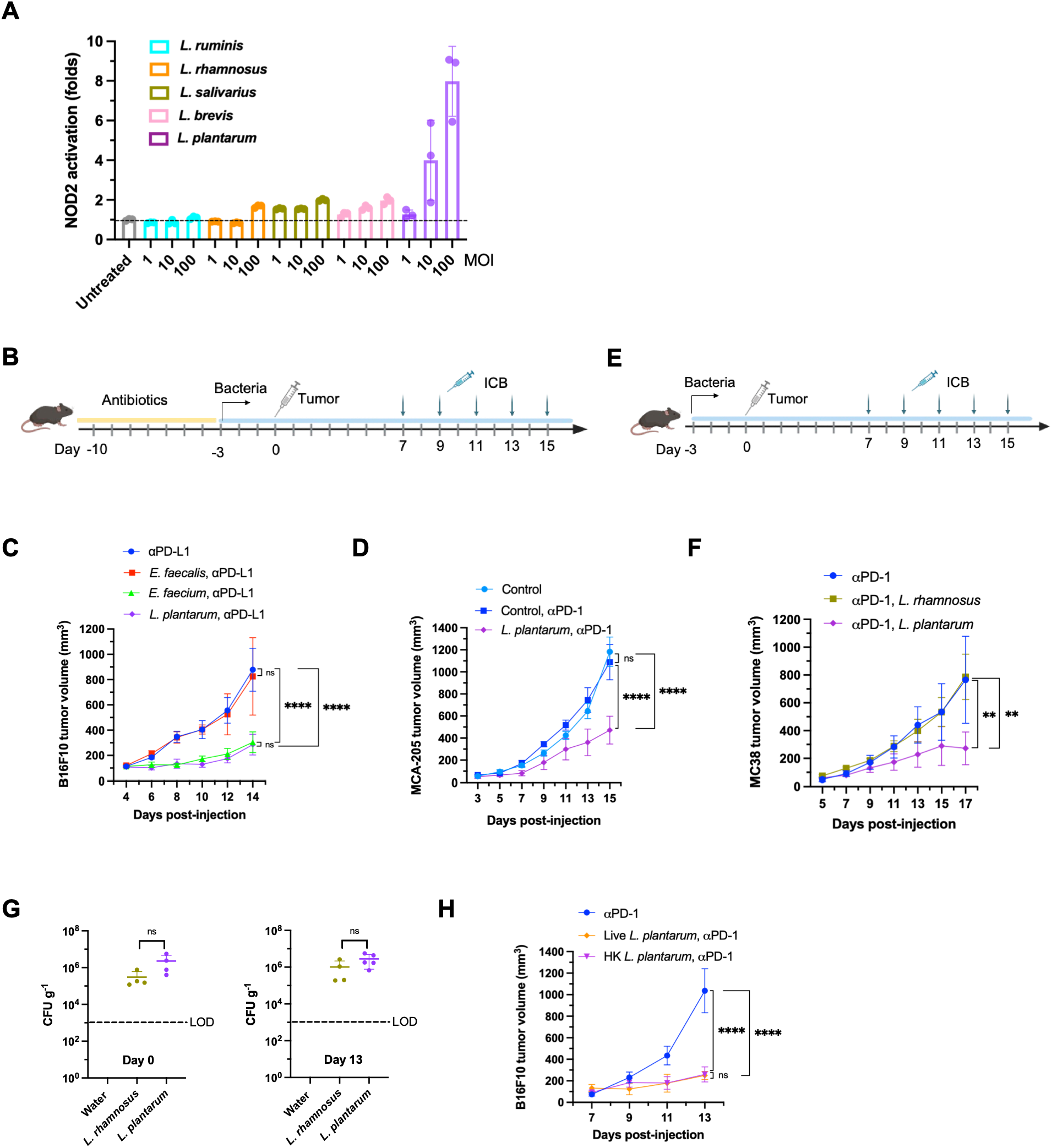
*L. plantarum* potentiates immune checkpoint blockade. **(A)** Activation of NOD2 by different *Lactobacillus* species. NOD2-HEKBlue reporter cells were stimulated with the indicated *Lactobacillus* species at varying multiplicity of infection (MOIs), corresponding to the indicated bacteria-to-cell ratios, and NF-κB activation was quantified and expressed relative to untreated control (n=3). Data are mean ± S.D. **(B)** Schematic of the antibiotic treatment and bacterial colonization regimen used in (C) and (D). Mice were treated with an antibiotic (Abx) cocktail for 1 week, followed by bacterial administration. Three days later, tumor cells were implanted subcutaneously. Immune checkpoint blockade (ICB) was initiated on day 7 after tumor implantation and administered intraperitoneally every other day thereafter. **(C)** B16F10 tumor growth in mice administered the indicated bacterial species. **(D)** MCA205 tumor growth in mice administered the indicated bacterial species. **(E)** Schematic of the bacterial supplementation regimen. Mice were supplemented with the indicated bacteria beginning 3 days before tumor implantation. ICB treatment was initiated on day 7 after tumor implantation. **(F)** MC38 tumor growth in mice supplemented with the indicated bacteria. **(G)** Fecal bacterial burdens were quantified by colony-forming unit (CFU) enumeration on days 0 and 13 after tumor implantation. **(H)** B16F10 tumor growth in mice orally gavaged with live or heat-killed *L. plantarum*.

We first tested *L. plantarum* in antibiotic-treated mice colonized with individual bacterial strains **(Fig. 1B)**. Similar to *E. faecium* but not *E. faecalis*, colonization with *L. plantarum* enhanced the response to ICB in both B16F10 melanoma **(Fig. 1C)** and MCA205 fibrosarcoma models **(Fig. 1D)**. We then asked whether this activity was retained in mice with an intact microbiota. In specific pathogen-free (SPF) mice, oral administration of *L. plantarum* but not *L. rhamnosus*, a weaker NOD2 stimulator, substantially reduced tumor burden when combined with ICB **(Fig. 1E and 1F)**.

This difference was not attributable to intestinal colonization, as both species achieved comparable colonization levels (**Fig. 1G**). Thus, the ability to enhance ICB is variable across *Lactobacillus* species with *L. plantarum* having a potent ICB antitumor activity, which is maintained in the context of a complex microbiota. We next investigated the bacterial features responsible for this activity. Heat-killed *L. plantarum* retained antitumor activity following either oral administration **(Fig. 1H)** or intratumoral delivery **(Fig. S1C)**, whereas cell-free culture supernatant failed to reproduce this effect. These findings suggested that the antitumor activity of *L. plantarum* is associated with bacterial cellular components rather than secreted metabolites and does not require bacterial viability. Because orally administered *L. reuteri* can translocate to tumors and locally promote antitumor immunity^17^, we asked whether *L. plantarum* similarly disseminated **(Fig. S1D)**. No viable bacteria were recovered from tumors of *L. plantarum*–supplemented mice, whereas spleen samples showed inconsistent CFU recovery **(Fig. S1E and S1F)**. These findings argue against direct tumor colonization as the basis for its antitumor activity.

Together, these results identify *L. plantarum* as a commensal bacterium that potentiates ICB through a bacterial cell-associated immunostimulatory activity without requiring bacterial viability or tumor translocation.

### NOD2 restrains *L. plantarum*-driven antitumor immunity

NOD2 is required for the antitumor activity of *E. faecium*^15,16^. Given the ability of *L. plantarum* to stimulate NOD2, we initially hypothesized that NOD2 might similarly mediate its antitumor activity. To test this, we compared the therapeutic responses to *E. faecium* and *L. plantarum* in *Nod2*-sufficient (*Nod2^+/−^*) and *Nod2*-deficient (*Nod2^-/-^*) littermates **(Fig. 2A)**. Consistent with our previous findings, *E. faecium*–mediated tumor control was lost in *Nod2^-/-^* mice. In contrast, *L. plantarum* retained its antitumor activity in the absence of NOD2, demonstrating that these two bacteria engage distinct host pathways to promote tumor control. Under this high-frequency oral gavage regimen, *L. plantarum* consistently showed a trend toward greater tumor control in *Nod2^-/-^* mice, which was also observed in the B16F10 tumor model (**Fig. S2A-S2C**). Flow cytometry analysis showed that *L. plantarum* increased intratumoral CD3⁺ T-cell accumulation in both genotypes, with a trend toward a greater increase in *Nod2^-/-^* mice **(Fig. S2D and S2E)**. Notably, increased accumulation of CD8⁺ T cells **(Fig. S2F)** and proliferating Ki-67⁺ CD8⁺ T cells **(Fig. S2G)** was observed only in *Nod2^-/-^* mice.

**Figure 2.**
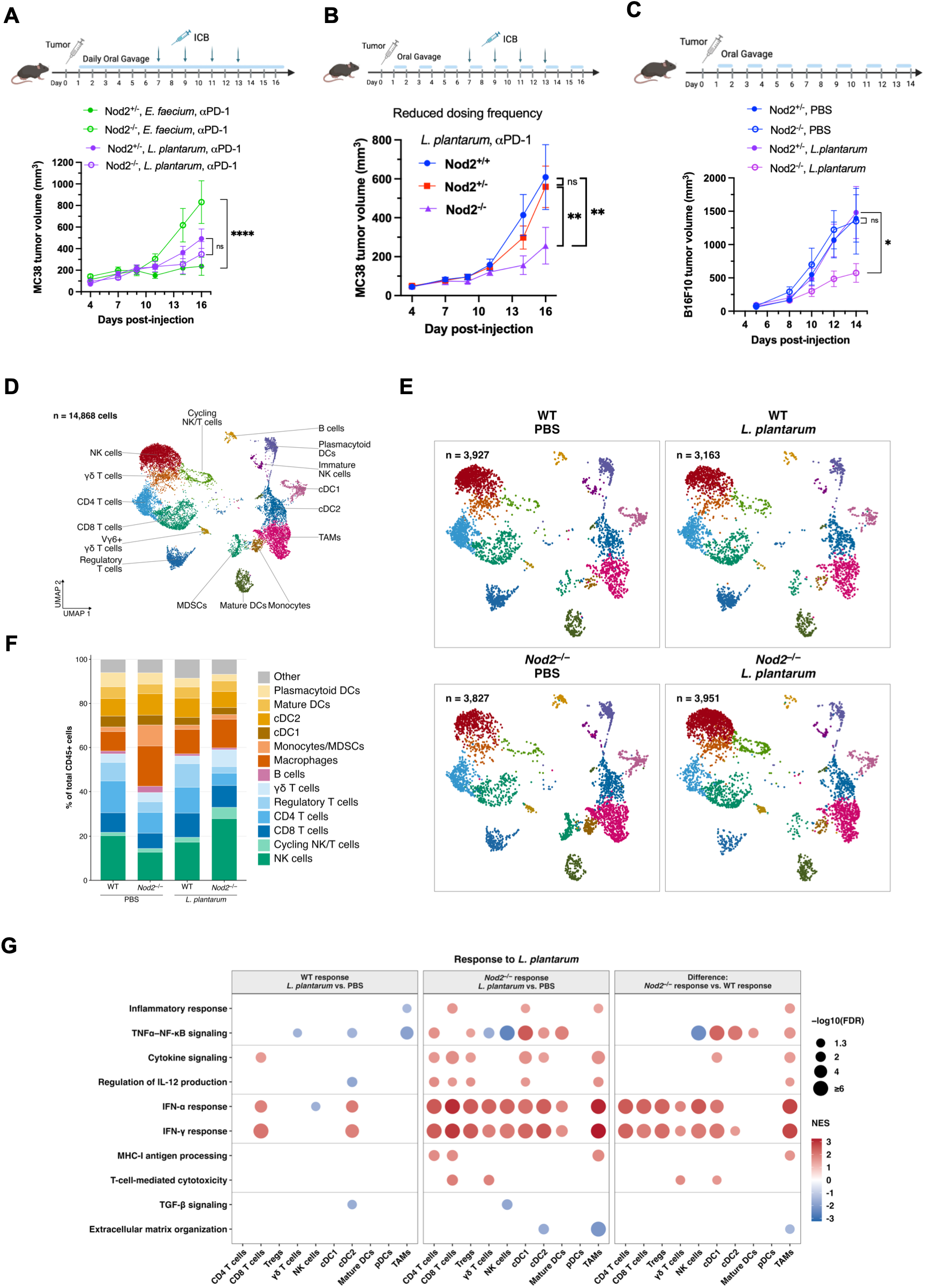
**NOD2 restrains *L. plantarum*–driven antitumor immunity and intratumoral interferon responses**. **(A)** Experimental schematic and MC38 tumor growth in *Nod2^+/-^*and *Nod2^-/-^* littermates treated with *L. plantarum* by daily oral gavage or *E. faecium* in the drinking water, together with anti–PD-1. Anti–PD-1 treatment was initiated on day 7 and administered every other day. The *E. faecium* groups were evaluated concurrently with the *L. plantarum* groups and were previously reported in^16^. **(B)** Experimental schematic and MC38 tumor growth in cohoused *WT*, *Nod2^+/-^*, and *Nod2^-/-^* mice receiving *L. plantarum* by oral gavage every other day beginning on day 1, together with anti–PD-1 initiated on day 7 and administered every other day. **(C)** Experimental schematic and B16F10 tumor growth in *Nod2^+/-^* and *Nod2^-/-^* littermates receiving PBS or *L. plantarum* by oral gavage every other day beginning on day 1 in the absence of ICB. **(D)** Uniform manifold approximation and projection (UMAP) of 14,868 tumor-infiltrating CD45^+^ immune cells isolated from WT and *Nod2^-/-^* mice treated with PBS or *L. plantarum* (n = 4 mice per group), colored by annotated cell type. **(E)** UMAPs from (E) split by experimental condition. The number of cells included in each condition is indicated. **(F)** Relative abundance of the indicated immune-cell populations, expressed as a percentage of total tumor-infiltrating CD45^+^ cells. Bars show the mean cell-type proportions across four mice per group. All annotated clusters were included in the analysis, and the biological replicate was the mouse. Differences in cell-type abundance were assessed using propeller/limma applied to logit-transformed per-mouse proportions, with two-sided testing and Benjamini–Hochberg correction within each contrast. **(G)** Cell type–resolved gene-set enrichment analysis of the *L. plantarum* response in WT mice, the *L. plantarum* response in *Nod2^-/-^* mice, and the genotype-by-treatment interaction. Enrichment analysis was performed on pseudobulk expression profiles using genes ranked by the limma-voom moderated *t* statistic. Only pathway–cell type combinations with FDR < 0.05 are shown. Bubble size represents −log10(FDR), capped at 6, and color indicates the normalized enrichment score (NES). Analyses were restricted to cell types represented by at least 10 cells in every mouse. Tumor-growth data are presented as mean ± SEM and were analyzed by a mixed-effects model with Tukey’s multiple comparisons post hoc test. Sample sizes are indicated in the figure or specified above. *\*P < 0.05; **P < 0.01; ****P < 0.0001; ns, not significant*.

We reasoned that the intensive dosing regimen might mask the inhibitory effect of NOD2 by providing sustained bacterial stimulation. We therefore reduced *L. plantarum* exposure by decreasing the frequency of administration. Under these conditions, *L. plantarum*-mediated tumor control was diminished in *Nod2*-sufficient (*Nod2^+/+^* and *Nod2^+/−^*) mice but remained evident in *Nod2^-/-^* mice **(Fig. 2B)**. Strikingly, this genotype-dependent difference was maintained even in the absence of ICB. *L. plantarum* had little effect on tumor growth in *Nod2*-sufficient mice but significantly suppressed tumor growth in *Nod2^-/-^* mice **(Fig. 2C)**. This difference was not attributable to altered intestinal bacterial abundance, as *L. plantarum* levels were comparable between genotypes **(Fig. S2H)**. Together, these findings identify NOD2 as a negative regulator of *L. plantarum*–mediated tumor control and prompted us to investigate how NOD2 shapes the intratumoral immune response to this bacterium.

### NOD2 restrains the *L. plantarum*–induced interferon program in the tumor immune microenvironment

To define how NOD2 deficiency alters the intratumoral immune response to *L. plantarum*, we performed single-cell RNA sequencing of 14,868 tumor-infiltrating CD45⁺ cells from PBS- or *L. plantarum*–treated WT and *Nod2^-/-^* mice (n = 4 mice per group). Unsupervised clustering identified 15 lymphoid and myeloid populations across all four conditions (**Fig. 2D and 2E**). Cell-composition analysis revealed that *L. plantarum* treatment in *Nod2^-/-^* mice markedly increased the representation of NK cells, from 12.8% to 28.0% of CD45⁺ cells, and altered the abundance of B cells, monocytes/MDSCs, cycling NK/T cells, γδ T cells, CD4⁺ T cells, and plasmacytoid dendritic cells (**Fig. 2F**). By contrast, no significant compositional changes were detected in WT tumors.

Cell-type-resolved pseudobulk analysis further demonstrated that *L. plantarum* induced stronger type I and type II interferon programs in *Nod2^-/-^* than in WT mice, particularly in tumor-associated macrophages and NK cells (**Fig. 2G)**. Enhanced TNFα–NF-κB signaling in dendritic cells and macrophages and increased T cell-mediated cytotoxicity were also observed, whereas extracellular matrix organization was reduced in tumor-associated macrophages. Notably, tumors from *Nod2^-/-^*mice exhibited lower interferon activity at baseline, indicating that the amplified response to *L. plantarum* was not attributable to elevated constitutive interferon signaling. Together, these findings indicate that NOD2 restrains *L. plantarum*–induced immune remodeling and interferon-associated antitumor responses in the tumor microenvironment.

### NOD2 suppresses *L. plantarum*-induced inflammatory responses *ex vivo*

In parallel with our *in vivo* studies, we asked whether NOD2 regulates the myeloid-cell response to *L. plantarum*. Bone marrow-derived macrophages (BMDMs) from *Nod2^+/−^* and *Nod2^-/-^*littermates were stimulated with *L. plantarum*, and a panel of cytokines and chemokines was measured **(Fig. S2I)**. In unstimulated BMDMs, mediator levels were generally low, and no clear genotype-dependent differences were detected. Following *L. plantarum* stimulation, however, CXCL1, CCL2, CCL5, and CXCL10 production was significantly greater in *Nod2^-/-^* BMDMs than in NOD2-sufficient controls, whereas the remaining detectable mediators did not differ significantly between genotypes (**Fig. S2J**). These results demonstrate that BMDMs derived from *Nod2^-/-^* mice exhibit enhanced chemokine responses to *L. plantarum*, consistent with NOD2 restraining *L. plantarum*–induced inflammatory responses.

### TLR2 is required for *L. plantarum*-driven antitumor immunity *in vivo*

Previous studies have implicated NOD2 as a negative regulator of TLR2-mediated responses^18,19^. TLR2 has also been implicated in the immunomodulatory and antitumor effects of *L. plantarum* strain IMB19 and its rhamnose-rich heteropolysaccharide^20^. However, whether host TLR2 is required for *L. plantarum*–mediated tumor control *in vivo* remained unknown. We first evaluated *L. plantarum* 8014-mediated tumor control in cohoused *Tlr2*-deficient (*Tlr2^-/-^* ) and *Tlr2*-sufficient (*Tlr2^+/-^* ) mice. Whereas *L. plantarum* inhibited tumor growth in TLR2-sufficient mice, this activity was lost in *Tlr2^-/-^* mice **(Fig. S3A)**. We subsequently confirmed the requirement for TLR2 using *Tlr2^+/-^* and *Tlr2^-/-^* littermates **(Fig. 3A–D)**. In the absence of bacterial treatment, tumors grew similarly in both genotypes **(Fig. 3B)**. Daily oral administration of *L. plantarum* in combination with anti-PD-L1 suppressed tumor growth in *Tlr2^+/-^* mice but failed to do so in *Tlr2^-/-^* mice **(Fig. 3C)**. In contrast, *E. faecium* in combination with anti-PD-L1 suppressed tumor growth in both genotypes **(Fig. 3D)**. Within each bacterial treatment group, fecal bacterial abundance was comparable between *Tlr2^+/-^* and *Tlr2^-/-^* mice **(Fig. S3B, C)**. Interestingly, in the absence of ICB, *E. faecium* lost tumor control activity in *Tlr2^+/-^* mice but exhibited marked antitumor activity in *Tlr2^-/-^* mice (**Fig. S3D)**. Thus, the requirement for TLR2 differs between bacterial species and treatment contexts, with TLR2 being required for *L. plantarum*–mediated tumor control but dispensable for, and under some conditions potentially restraining, the antitumor activity associated with *E. faecium*.

**Figure 3.**
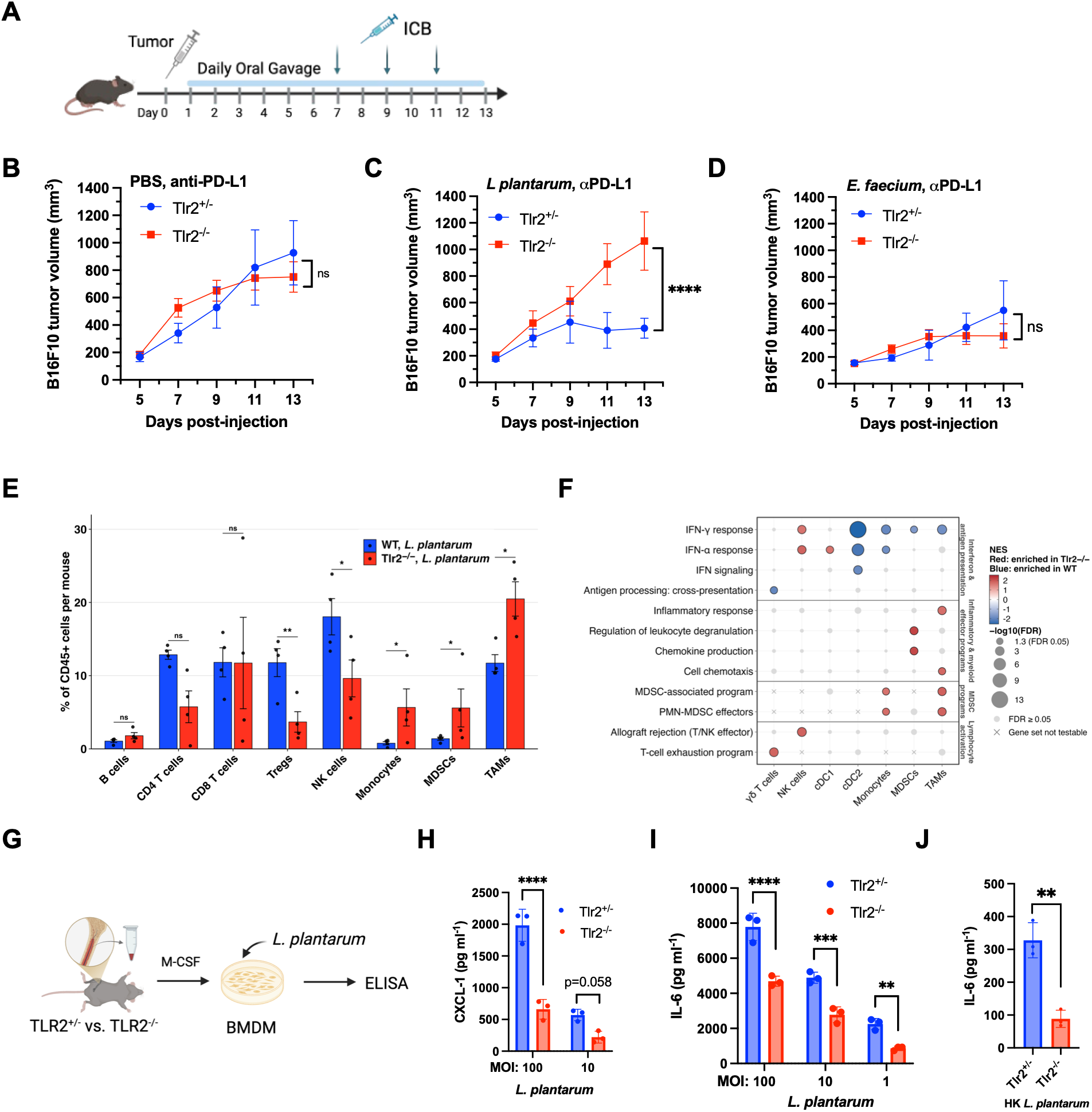
**TLR2 is required for *L. plantarum*–driven antitumor immunity**. **(A)** Experimental schematic. *Tlr2^+/-^* and *Tlr2^-/-^* littermates bearing B16F10 tumors received PBS or *L. plantarum* by daily oral gavage, or *E. faecium* in the drinking water, in combination with anti–PD-L1. Anti–PD-L1 treatment was initiated on day 7 and administered every other day. **(B–D)** B16F10 tumor growth in mice treated with PBS **(B)**, *L. plantarum* **(C)**, or *E. faecium* **(D)**. **(E)** Mouse-level proportions of eight major tumor-infiltrating immune populations, expressed as a percentage of total CD45+ cells in *L. plantarum*–treated WT and *Tlr2^-/-^* mice. Bars indicate group means, error bars indicate SEM, and points represent individual mice (n = 4 mice per genotype). Raw two-sided *P* values were calculated using propeller with limma-moderated *t* tests applied to logit-transformed per-mouse proportions. Benjamini–Hochberg-adjusted *q* values were calculated across all 15 annotated immune populations. Results for all 15 annotated populations are shown in Figure S3F. **(F)** Cell type–resolved gene-set enrichment analysis of tumor-infiltrating immune cells from *L. plantarum*–treated WT and *Tlr2^-/-^* mice. **(G)** Schematic of the *ex vivo* macrophage-stimulation experiment. Bone marrow cells isolated from *Tlr2^+/-^*and *Tlr2^-/-^*littermates were differentiated with M-CSF for 7 days. The resulting BMDMs were stimulated overnight with *L. plantarum*, after which culture supernatants were collected for ELISA. **(H–J)** Cytokine concentrations in culture supernatants from *Tlr2^+/-^* and *Tlr2^-/-^* BMDMs stimulated overnight with *L. plantarum*. CXCL1 **(H)** and IL-6 **(I)** concentrations following stimulation with live *L. plantarum* at the indicated multiplicities of infection (MOIs), and IL-6 concentrations following stimulation with heat-killed (HK) *L. plantarum* **(J)**. Tumor-growth data are presented as mean ± SEM and were analyzed by a mixed-effects model with Tukey’s multiple comparisons post hoc test. *Ex vivo* data are presented as mean ± SEM from 3 independent biological replicates and were analyzed using one-way ANOVA followed by Tukey’s multiple-comparisons test in **(H)** and **(I)** and a two-sided unpaired *t* test in **(J)**. *\*P < 0.05; **P < 0.01; ***P < 0.001; ****P < 0.0001; ns, not significant*.

### Loss of TLR2-dependent tumor control is associated with altered antitumor immune programs

Given the loss of *L. plantarum*–mediated tumor control in *Tlr2^-/-^* mice, we next asked how TLR2 deficiency altered the tumor immune response to *L. plantarum*. We performed scRNA sequencing of tumor-infiltrating CD45^+^ immune cells from WT and *Tlr2^-/-^* mice following *L. plantarum* treatment. TLR2 deficiency was associated with broad changes in the tumor immune compartment, including reduced representation of Tregs and NK cells and increased representation of monocytes, MDSCs, and tumor-associated macrophages **(Fig. 3E, S3E and S3F)**. Cell type-resolved pseudobulk analysis revealed stronger IFN-α- and IFN-γ-response signatures in WT cDC2 and monocytes, together with stronger IFN-γ-response signatures in WT MDSCs and TAMs **(Fig. 3F)**. Conversely, *Tlr2⁻/⁻* mice exhibited enrichment of chemokine-production and leukocyte-degranulation programs in MDSCs, inflammatory and chemotactic programs in TAMs, and MDSC-associated programs in monocytes and TAMs. Together, these findings associate the loss of *L. plantarum*–mediated tumor control in TLR2-deficient mice with increased representation of myeloid populations and reduced interferon-response programs in several myeloid compartments.

### TLR2 promotes *L. plantarum*–induced inflammatory responses *ex vivo*

Because the tumor immune landscape reflects both direct bacterial sensing and secondary changes within the tumor microenvironment, we next examined the requirement for TLR2 in the direct macrophage response to *L. plantarum ex vivo*. BMDMs from *Tlr2^+/-^* and *Tlr2^-/-^* littermates were stimulated with *L. plantarum* and analyzed by ELISA. In contrast to the enhanced responses observed upon NOD2 deficiency, loss of TLR2 markedly reduced *L. plantarum*–induced CXCL1 **(Fig. 3H)** and IL-6 production **(Fig. 3I)**. IL-6 production induced by heat-killed *L. plantarum* was similarly diminished in *Tlr2^-/-^* BMDMs **(Fig. 3J)**, consistent with TLR2-dependent recognition of a heat-stable, cell-associated bacterial component. Together, these findings establish a requirement for TLR2 in the acute macrophage response to *L. plantarum*, while distinguishing this direct sensing response from the broader immune remodeling observed in TLR2-deficient tumors.

### CPS is dispensable for *L. plantarum*–induced TLR2 activation

We next sought to identify the *L. plantarum* molecular factors responsible for TLR2 activation. Several microbe-associated molecular patterns, including lipoproteins and lipopeptides, lipoteichoic acid, and capsular polysaccharides (CPS), have been implicated in TLR2 signaling^21^. We initially focused on CPS because a rhamnose-rich heteropolysaccharide purified from *L. plantarum* IMB19 was previously proposed to activate TLR2 directly and enhance ICB^20^. However, this activity was not genetically validated using an isogenic bacterial mutant. Conversely, deletion of *cps* from *L. plantarum* WCFS1 (the commonly studied strain) was reported to increase TLR2 reporter activity^22^, suggesting that the high molecular weight CPS may instead mask underlying TLR2 ligands and limit their recognition.

To assess whether differences in CPS biosynthetic capacity could account for strain-dependent immune stimulation and antitumor activity, we compared CPS gene clusters of *L. plantarum* IMB19, WCFS1, and ATCC 8014. Comparative genomic analysis showed that only the *cps4* cluster and the complete rhamnose-biosynthesis gene set (*rmlA–D*) were conserved across all three strains, whereas WCFS1 contained three additional *cps* clusters **(Fig. S4A)**. Consistent with *L. plantarum* WCFS1 strain containing additional CPS gene clusters (cps1, cps2, cps3) that are absent in *L. plantarum* 8014 strain, crude capsular polysaccharide profiling revealed more high molecular weight CPS structures in WCFS1 than in ATCC 8014 **(Fig. S4B)**. To further characterize immunogenicity of these CPS-variable *L. plantarum* strains, we evaluated their antitumor activity *in vivo* and compared their immunostimulatory effects *in vitro*. Unlike live 8014, live WCFS1 failed to enhance ICB-mediated tumor control **(Fig. S4C)**. However,, HK WCFS1 displayed a comparative antitumor activity to HK 8014, suggesting that heat treatment exposes or releases a cell-associated immunostimulatory component **(Fig. S4D)**. Despite their different antitumor activities *in vivo*, WCFS1 and 8014 elicited comparable NOD2 and TLR2 reporter activation when tested as either live or HK bacteria **(Fig. S4E)**. The two strains also induced similar CXCL10 and IL-6 production in BMDMs and comparable NF-κB and IRF reporter responses in human THP-1 cells **(Fig. S4F–S4I)**. These findings indicate that differences in CPS abundance or organization do not solely explain the ability of these strains to stimulate innate immune signaling *in vitro* and suggest that the reduced activity of live WCFS1 *in vivo* may instead reflect differential accessibility of an underlying cell-associated factor.

To test the contribution of CPS more directly, we established a gene editing approach for *L. plantarum* 8014 to be able to make isogenic mutants. This allowed us to generate a panel of isogenic 8014 mutants targeting key enzymes in CPS biosynthesis. Deletion of *wzx* in two different loci, which encodes a putative polysaccharide repeat-unit flippase, produced no detectable change in the polysaccharide profile, potentially reflecting functional redundancy among CPS loci **(Fig. S5A)**. In contrast, deletion of another CPS synthesis gene rmlX, abolished detectable CPS production **(Fig. S5A)**. Nevertheless, neither deletion altered NOD2 or TLR2 reporter activation by *L. plantarum* 8014 **(Fig. S5B and S5C)**. Together, these studies with isogenic CPS mutants indicate that CPS are dispensable for *L. plantarum*–induced TLR2 activation. We therefore shifted our attention to bacterial lipoproteins, which are potent TLR2 ligands, as an alternative source of molecular mediators of *L. plantarum* TLR2-dependent activity.

### Lipoprotein drives TLR2 activation and contributes to *L. plantarum*–mediated antitumor immunity

Because disruption of CPS biosynthesis did not diminish TLR2 activation of *L. plantarum*, we next examined whether bacterial lipoproteins mediate this activity. Lipoproteins are potent TLR2 ligands whose immunostimulatory activity depends on lipid modification by lipoprotein diacylglyceryl transferase (*lgt*), conserved across *L. plantarum* strains. Moreover, LPPs induced-TLR2 signaling from different bacteria have been shown to be important for cancer immunotherapy outcomes^26,27^. We therefore generated an *L. plantarum lgt* mutant and functionally characterized it. Using click chemistry-based profiling of bacterial LPPs metabolically labeled by alkynyl-fatty acids^28^, we observed a marked loss of lipidated proteins in the *lgt* mutant **(Fig. S6A)**, whereas bacterial growth was not significantly affected **(Fig. S6B)**. We also performed peptidoglycan profiling and found that the lipoprotein mutant did not alter muropeptide composition, which was also conserved across different strains **(Fig. S6C and D)**. Consistently, the *lgt* mutant did not alter the activation of NOD2 in reporter cell assay **(Fig. S6E)**, but completely abolished *L. plantarum*–induced TLR2/6 reporter activation **(Fig. 4A).** These findings identify LPPs as major mediators of TLR2 activation by *L. plantarum* 8014.

**Figure 4.**
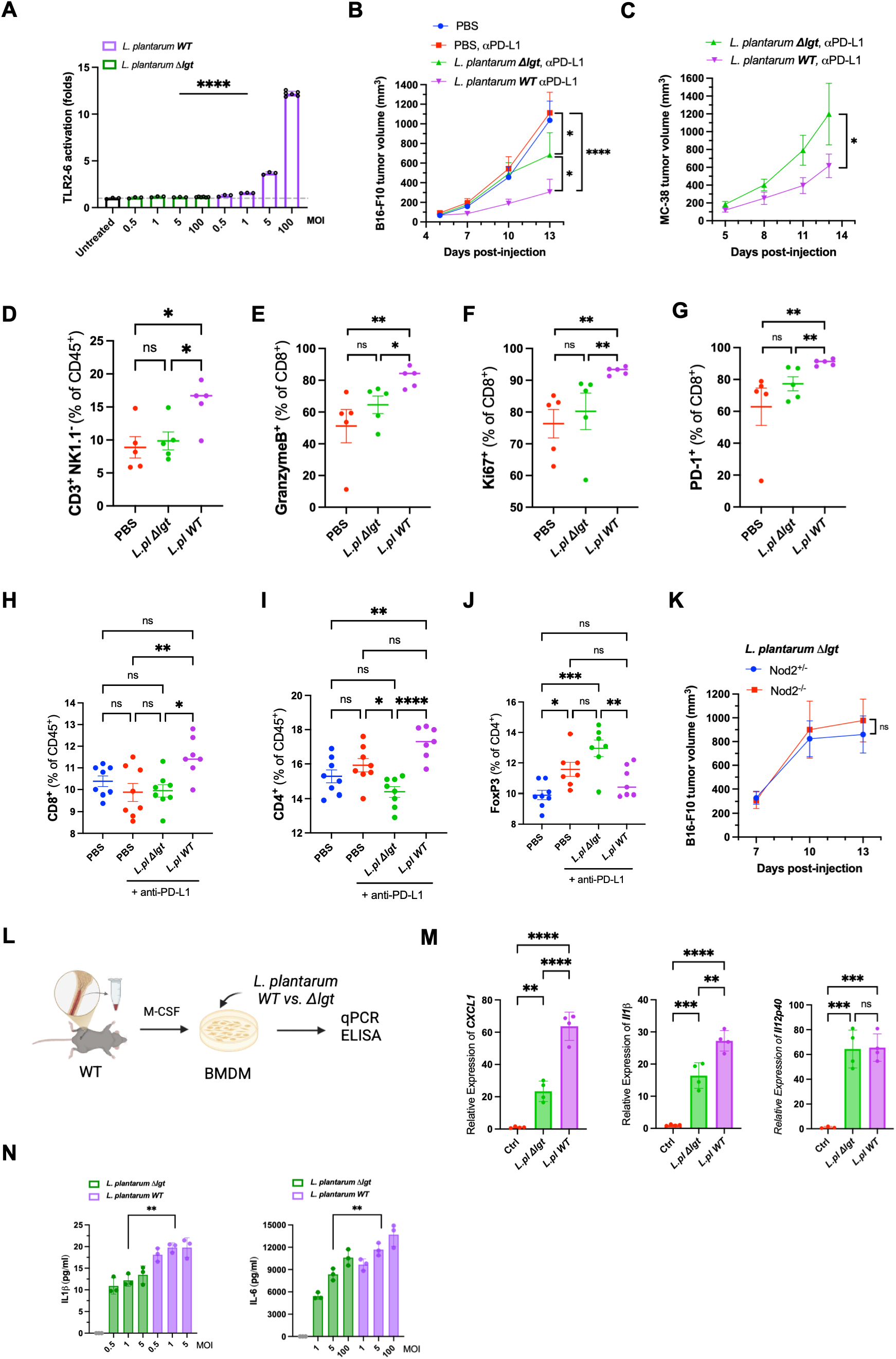
*L. plantarum* lipoproteins promote TLR2-dependent antitumor immunity. **(A)** TLR2/6 reporter activation by *L. plantarum* ATCC 8014 WT and Δ*lgt* at the indicated MOIs. Reporter activity is shown as fold activation relative to untreated cells. **(B)** B16F10 tumor growth in mice receiving PBS, PBS plus anti–PD-L1, *L. plantarum* Δ*lgt* plus anti–PD-L1, or *L. plantarum* WT plus anti–PD-L1. Bacteria were administered daily by oral gavage. **(C)** MC38 tumor growth in mice receiving daily oral administration of *L. plantarum* WT or Δ*lgt* in combination with anti–PD-L1. **(D–G)** Flow-cytometric quantification of tumor-infiltrating **(D)** CD3⁺NK1.1⁻ T cells as a percentage of CD45⁺ cells, **(E)** Granzyme B⁺ CD8⁺ T cells, **(F)** Ki-67⁺ CD8⁺ T cells, and **(G)** PD-1⁺ CD8⁺ T cells from the B16F10 tumors. **(H–J)** Flow-cytometric quantification of splenic **(H)** CD8⁺ T cells as a percentage of CD45⁺ cells, **(I)** CD4⁺ T cells as a percentage of CD45⁺ cells, and **(J)** regulatory T cells as a percentage of CD4⁺ T cells from the indicated treatment groups. **(K)** B16F10 tumor growth in *Nod2^+/-^*and *Nod2^-/-^* littermates receiving daily oral administration of *L. plantarum* Δ*lgt*. **(L)** Schematic of the *ex vivo* macrophage-stimulation experiment. Bone marrow cells from WT mice were differentiated with M-CSF for 7 days, and the resulting BMDMs were stimulated with *L. plantarum* WT or Δ*lgt* before analysis by qPCR and ELISA. **(M)** Relative expression of *Cxcl1*, *Il1β*, and *Il12b* in BMDMs left untreated or stimulated with *L. plantarum* WT or Δ*lgt*, as determined by qPCR. **(N)** IL-1β and IL-6 concentrations in culture supernatants from BMDMs stimulated with *L. plantarum* WT or Δ*lgt* at the indicated MOIs. Tumor-growth data in **(B)**, **(C)**, and **(K)** are presented as mean ± SEM and were analyzed using a mixed-effects model followed by Tukey’s multiple-comparisons test. Flow-cytometry data are presented as mean ± SEM; each symbol represents an individual mouse. Statistical comparisons in **(D–J)** were performed using one-way ANOVA followed by Tukey’s multiple-comparisons test. *Ex vivo* data are presented as mean ± SEM from 3 independent biological replicates and were analyzed using one-way ANOVA followed by Tukey’s multiple-comparisons test. *\*P < 0.05; **P < 0.01; ***P < 0.001; ****P < 0.0001; ns, not significant*.

We next tested whether lipoproteins contribute to the antitumor activity of *L. plantarum in vivo*. Tumor-bearing mice received daily oral gavage with WT *L. plantarum* or the *lgt* mutant in combination with ICB. Compared with WT *L. plantarum*, the *lgt* mutant exhibited a reduced ability to control tumor growth in both the B16F10 and MC38 tumor models **(Fig. 4B and Fig. 4C)**. Consistent with its impaired therapeutic activity, the *lgt* mutant failed to promote the intratumoral accumulation of CD3⁺ T cells observed with WT *L. plantarum* **(Fig. 4D)** and elicited weaker CD8⁺ T-cell responses, as reflected by reduced granzyme B, Ki-67, and PD-1 expression **(Fig. 4E–4G)**. WT *L. plantarum* also increased splenic CD4⁺ and CD8⁺ T-cell responses, whereas these effects were not observed with the *lgt* mutant **(Fig. 4H and 4I)**. Moreover, splenic CD4⁺ and CD8⁺ T-cell abundance negatively correlated with tumor size in mice treated with WT *L. plantarum*, but not in those treated with the *lgt* mutant **(Fig. S6F)**, linking lipoprotein-dependent systemic T-cell responses to improved tumor control. Consistent with its impaired induction of systemic effector T-cell responses, the *lgt* mutant increased the proportion of splenic Foxp3⁺ CD4⁺ regulatory T cells relative to WT *L. plantarum* **(Fig. 4J)**, suggesting a shift toward a more regulatory systemic immune state. Together, these findings indicate that LPPs promote intratumoral and systemic effector T-cell responses while limiting systemic regulatory T-cell accumulation, thereby contributing to *L. plantarum*–mediated tumor control.

### *L. plantarum* lipoproteins are required for enhanced tumor control in NOD2-deficient mice

We next asked whether the enhanced antitumor response to *L. plantarum* in *Nod2^-/-^* mice required Lgt-dependent TLR2 stimulation. Whereas WT *L. plantarum* exhibited robust antitumor activity in *Nod2^-/-^* mice **(Fig. 2B and 2C)**, the *lgt* mutant failed to suppress tumor growth, even in the *Nod2^-/-^* mice **(Fig. 4K)**. These findings indicate that loss of Lgt-dependent TLR2 immune stimulation abolishes the enhanced response to *L. plantarum* associated with NOD2 deficiency, further supporting the dependence of this phenotype on TLR2 activation.

### Lgt-dependent lipoproteins promote *L. plantarum*–induced inflammatory responses *ex vivo*

In parallel with our *in vivo* studies, we asked whether Lgt-dependent lipoproteins contribute to the macrophage inflammatory response to *L. plantarum*. BMDMs were stimulated with WT *L. plantarum* or the *lgt* mutant, and inflammatory gene expression and cytokine production were measured **(Fig. 4L)**. Compared with WT *L. plantarum*, the *lgt* mutant induced significantly lower expression of *Cxcl1* and *Il1β*, whereas *Il12b* expression was unchanged **(Fig. 4M)**. Consistent with these transcriptional changes, the *lgt* mutant induced less IL-1β and IL-6 **(Fig. 4N)** production than WT *L. plantarum*. These findings indicate that Lgt-dependent lipoproteins selectively promote *L. plantarum*–induced inflammatory responses in macrophages.

### NOD2 deficiency enhances TLR2 agonist-induced inflammatory responses in mouse and human myeloid cells

To determine whether the inhibitory effect of NOD2 extends beyond bacterial stimulation, we examined responses to the defined TLR2 agonist Pam3CSK4, which is a synthetic triacylated lipopeptide that mimics bacterial lipoproteins. BMDMs from WT, *Nod2^-/-^*, and *Tlr2^-/-^* mice were left untreated or stimulated with Pam3CSK4 and analyzed by bulk RNA sequencing and ELISA **(Fig. 5A)**. WT and *Nod2^-/-^* macrophages mounted similarly broad transcriptional responses to Pam3CSK4, whereas the response was nearly abolished in *Tlr2^-/-^* cells **(Fig. S7A)**. Direct comparison of Pam3CSK4-stimulated *Nod2^-/^* and WT macrophages revealed selective enhancement of an immune-effector gene module rather than uniform amplification of the entire TLR2 response. This module included the interferon-inducible GTPases *Gbp2*, *Gbp3*, and *Gbp7*, the Th1-associated cytokine genes *Il12b* and *Il15*, and the antimicrobial effector gene *Nos2* **(Fig. 5B)**. Increased expression of *Ifnar1* and *Stat2* further suggested enhanced type I interferon responsiveness in the absence of NOD2, although the functional contribution of this pathway remains to be established. At the protein level, Pam3CSK4 induced greater production of CXCL10, CCL2, CCL5, and CXCL1 in *Nod2^-/-^* BMDMs than in NOD2-sufficient controls **(Fig. 5C)**. This pattern mirrored the enhanced chemokine response elicited by *L. plantarum* in the absence of NOD2. In contrast, Pam3CSK4-induced IL-6 and TNF-α production was comparable between the two genotypes, indicating that NOD2 deficiency selectively amplifies a chemokine-dominant response rather than uniformly increasing all TLR2-dependent inflammatory outputs. These responses were diminished in *Tlr2^-/-^* BMDMs, confirming their dependence on TLR2 signaling. To determine whether this regulatory relationship extends to human myeloid cells, we generated an *NOD2*-deficient human THP-1 NF-κB reporter cell line and compared its response with that of the parental control. Pam3CSK4 induced dose-dependent NF-κB activation in both cell lines, but responses were consistently greater in *NOD2*-deficient cells across the concentration range tested **(Fig. 5D)**. These findings extend our observations from primary mouse macrophages to a human myeloid-cell model and indicate that NOD2-mediated restraint of TLR2-driven inflammatory signaling is observed across species.

**Figure 5.**
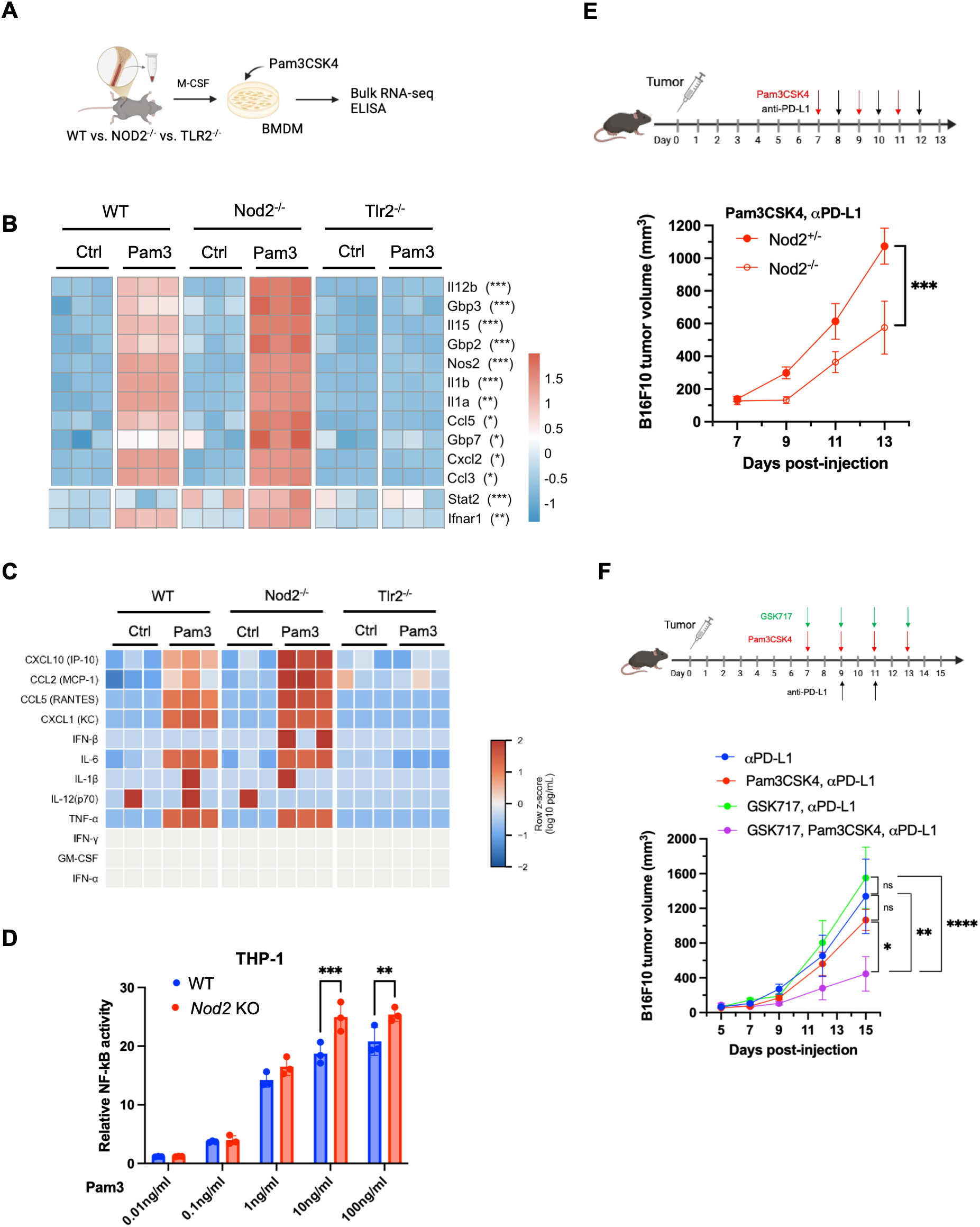
Genetic or pharmacological inhibition of NOD2 enhances TLR2 agonist–driven antitumor immunity. **(A)** Schematic of the ex vivo macrophage-stimulation experiment. Bone marrow cells from WT, *Nod2^-/-^*, and *Tlr2^-/-^*mice were differentiated with M-CSF for 7 days. The resulting BMDMs were left untreated or stimulated with the TLR2/1 agonist Pam3CSK4 before bulk RNA sequencing and cytokine analysis. **(B)** Expression of selected inflammatory, interferon-associated, and T-cell-polarizing genes in WT, *Nod2^-/-^*, and *Tlr2^-/-^* BMDMs left untreated or stimulated with Pam3CSK4. Expression values are shown as row-scaled normalized expression. Statistical significance refers to the comparison between Pam3CSK4-treated *Nod2^-/-^* and WT BMDMs. **(C)** Multiplex analysis of cytokines and chemokines in culture supernatants from WT, *Nod2^-/-^*, and *Tlr2^-/-^* BMDMs left untreated or stimulated with Pam3CSK4. Values are shown as row-scaled relative concentrations. **(D)** Relative NF-κB reporter activity in WT and NOD2-deficient THP-1 cells stimulated with the indicated concentrations of Pam3CSK4. **(E)** Schematic and B16F10 tumor growth in *Nod2^+/-^* and *Nod2^-/-^* littermates receiving Pam3CSK4 and anti–PD-L1 by intraperitoneal injection on the indicated days. **(F)** Schematic and B16F10 tumor growth in WT specific pathogen-free mice receiving anti–PD-L1 alone or in combination with Pam3CSK4, the NOD2 inhibitor GSK717, or both compounds. GSK717 was administered by intraperitoneal injection 1 h before Pam3CSK4 on the indicated treatment days. Tumor-growth data in **(E)** and **(F)** are presented as mean ± SEM and were analyzed using a mixed-effects model followed by Tukey’s multiple-comparisons test. Data in **(D)** are presented as mean ± SEM; each symbol represents an independent biological replicate. Statistical comparisons were performed using two-way ANOVA followed by Šídák’s multiple-comparisons test. *\*P < 0.05; **P < 0.01; ***P < 0.001; ****P < 0.0001; ns, not significant*.

### Genetic or pharmacological disruption of NOD2 potentiates TLR2 agonist-mediated tumor control

TLR2 agonists have been explored as cancer immunotherapy adjuvants. Previous work showed that intratumoral administration of Pam3CSK4 enhanced the efficacy of CTLA-4 blockade^29^; however, whether systemic Pam3CSK4 administration can potentiate other forms of ICB, and whether its activity is influenced by host NOD2 signaling, remained unclear. We therefore tested whether NOD2 deficiency enhances the antitumor activity of systemically administered Pam3CSK4. Tumor-bearing *Nod2^+/-^* and *Nod2^-/-^* littermates received intraperitoneal Pam3CSK4 together with ICB. Pam3CSK4 produced significantly greater tumor control in *Nod2^-/-^* mice than in *Nod2^+/-^* controls **(Fig. 5E)**, demonstrating that loss of NOD2 enhances the therapeutic response to systemic TLR2 stimulation.

We next asked whether pharmacological inhibition of NOD2 signaling could similarly enhance responses to TLR2 agonists. GSK717 is a reported inhibitor of muramyl dipeptide (MDP)-induced NOD2 signaling^30^. In RAW 264.7 reporter cells, GSK717 suppressed NF-κB activation induced by MDP while modestly enhancing responses to Pam2CSK4 and Pam3CSK4 **(Fig. S7B)**. We therefore asked whether GSK717 could enhance the antitumor activity of Pam3CSK4 *in vivo*. Tumor-bearing mice receiving ICB were treated with GSK717 one hour before each intraperitoneal administration of Pam3CSK4. Neither GSK717 nor Pam3CSK4 alone produced substantial tumor control under these conditions, whereas their combination significantly reduced tumor growth compared with Pam3CSK4 alone **(Fig. 5F and S7C)**. Body weight remained stable across the treatment groups **(Fig. S7D)**, providing no evidence of overt treatment-associated toxicity. Together, these findings provide genetic and pharmacological evidence that attenuation of NOD2 signaling can potentiate TLR2 agonist-mediated tumor control.

Finally, we asked whether prior NOD2 stimulation reciprocally attenuates the acute inflammatory response to Pam3CSK4 *in vivo*. Intraperitoneal administration of Pam3CSK4 induced robust circulating CXCL1, whereas pretreatment with the NOD2 agonist MDP markedly diminished this response **(Fig. S7E)**. Thus, pharmacological activation of NOD2 was associated with suppression of Pam3CSK4-induced inflammation, providing reciprocal support for a model in which NOD2 restrains TLR2-driven inflammatory responses *in vivo*.

## Discussion

In this study, we found that the antitumor activity of a commensal bacterium depends on interactions between host microbial-sensing pathways. *L. plantarum* enhanced ICB efficacy even after heat inactivation. Its antitumor activity required host TLR2 and was diminished by disruption of bacterial LPPs, whereas NOD2 deficiency enhanced tumor control. This contrasts with *E. faecium*, whose antitumor activity requires NOD2, demonstrating that the same receptor can support responses to one commensal while restraining those to another. Genetic loss or pharmacological inhibition of NOD2 also enhanced the antitumor response to the synthetic TLR2 agonist Pam3CSK4. Together, these results suggest that the therapeutic activity of bacteria and microbial products depends on both the pathways they activate and the host signals that modulate those responses. Therefore, the characterization of specific ICB-promoting microbiota and probiotic species and the elucidation of their mechanisms of action is crucial for their translation towards cancer immunotherapy.

*Lactobacilli* are phylogenetically and functionally diverse^31^, and their reported antitumor effects arise through distinct mechanisms^5^. For example, an exopolysaccharide from *L. delbrueckii* subsp. *bulgaricus* induced CCR6⁺ CD8⁺ T cells in Peyer’s patches and enhanced ICB efficacy against CCL20-expressing tumors^32^. A different bacterial product, *L. paracasei*-derived phytosphingosine increased tumor-cell MHC class I expression through an MYD88–NF-κB–NLRC5 pathway^33^. Other effects depend on microbial metabolism. *L. reuteri* converted dietary tryptophan to indole-3-aldehyde, which promoted AhR-dependent CD8⁺ T cell responses and improved ICB efficacy in melanoma^17^. In a distinct setting, *L. reuteri*-derived reuterin suppressed colorectal tumor growth by altering cancer-cell redox balance and protein translation^34^. Tryptophan metabolism by other lactobacilli can also influence ICB responses. *L. johnsonii* cooperates with *Clostridium sporogenes* to produce indole-3-propionic acid, which supports progenitor-exhausted CD8⁺ T cells and enhances ICB efficacy^35^, By contrast, *L. gallinarum*-derived indole-3-lactic acid suppresses colorectal tumorigenesis and promotes apoptosis of colorectal cancer cells^36^. Together, these studies illustrate that related bacteria can influence cancer through different metabolites and host processes. Despite all these studies and others, the precise mechanisms of action for many probiotic species/strains *in vivo* are still unclear, due to limited analysis of isogenic mutant strains, which restricts the development and prediction of precise/effective probiotics for cancer immunotherapy in specific patient populations.

Functional differences are also evident among *L. plantarum* strains. For example, L168-derived indole-3-lactic acid was shown to increase *Il12a* expression in dendritic cells, prime CD8⁺ T cell immunity, and reduce colorectal tumorigenesis^37^. Outside cancer, strain WJL supported growth during chronic undernutrition in mice^38^, an effect subsequently shown to require NOD2 sensing by intestinal epithelial cells^39^. In another tumor model, IMB19 promoted antitumor immunity through a capsular heteropolysaccharide that activates TLR2 and induces iron sequestration by tumor-associated macrophages^20^. Our findings identify an additional layer of regulation: NOD2 restrains the TLR2-dependent antitumor response to 8014, and disruption of bacterial lipoproteins diminished this activity. We also observed differences between *L. plantarum* strains. Live 8014 suppressed tumor growth, whereas live WCFS1 did not, despite similar receptor activation *ex vivo*. Heat-killed WCFS1 gained antitumor activity, raising the possibility that heat treatment increases the accessibility of immunostimulatory components. Whether surface polysaccharides account for this difference remains to be determined. Together, these findings suggest that both bacterial components and host context shape the activity of *L. plantarum*, and that *ex vivo* receptor activation alone may not predict antitumor activity *in vivo*.

NOD2 has previously been shown to limit TLR2-mediated inflammatory responses^18,19,40^, and protect against inflammation-associated colorectal tumorigenesis^41^. Our findings extend this regulatory relationship to microbiota-driven antitumor immunity. NOD2 deficiency enhanced chemokine and interferon-associated responses to *L. plantarum* and Pam3CSK4, while genetic loss or pharmacological inhibition of NOD2 improved the antitumor response to TLR2 stimulation. Previous studies showed that MDP–NOD2 stimulation induces IRF4-dependent suppression of subsequent responses to multiple TLR ligands^42^. IRF4 can inhibit TLR signaling by competing with IRF5 for MyD88 binding^43^. Following NOD2 activation, IRF4 also inhibits K63-linked polyubiquitination of RIPK2 and TRAF6, thereby limiting downstream NF-κB activation^44^. However, we did not detect increased *Irf4* transcript or IRF4 protein abundance following *L. plantarum* stimulation under the conditions examined. ATG16L1-dependent autophagy has also been shown to restrict TLR- and NLR-mediated inflammatory signaling^45^, and may contribute to the regulation observed here. Whether IRF4, autophagy, or other mechanisms underlie NOD2-mediated restraint remains to be determined. While our studies show stronger TLR2 responses in the absence of NOD2, further studies are needed to define the mechanisms on how NOD2 negatively regulates TLR2-dependent responses to *L. plantarum*.

TLR2 agonists have shown antitumor activity in several preclinical settings, although their efficacy depends on the treatment context. Intratumoral Pam3CSK4 administration enhanced anti–CTLA-4 efficacy by increasing FcγRIV expression on tumor-associated macrophages and promoting depletion of intratumoral regulatory T cells^29^. The small-molecule TLR1/2 agonist Diprovocim also enhanced anti–PD-L1 efficacy when administered intramuscularly with a tumor-associated antigen^46^. TLR2 activation can additionally modify the response to other innate agonists: TLR2 preactivation limited STING-induced PD-L1 expression and protumoral functions in tumor-infiltrating monocytes, improving STING agonist efficacy^47^. Our findings identify NOD2 as another determinant of the therapeutic response to TLR2 stimulation in the context of host-microbiota cancer immunotherapy. Under our experimental conditions, intraperitoneal Pam3CSK4 did not improve tumor control in NOD2-sufficient mice, whereas genetic loss or pharmacological inhibition of NOD2 enhanced its activity in combination with ICB. Thus, limited efficacy of a TLR2 agonist can reflect restraint imposed by another microbial-sensing pathway such as NOD2. The response to Pam3CSK4 combined with GSK717 provides a proof of concept for pairing innate receptor activation with inhibition of a pathway that constrains its antitumor effects.

Our findings also have implications for the clinical use of microbiota-based therapies. A recent study associated inherited NOD2 loss-of-function variants with favorable responses to cancer immunotherapy, including anti–PD-1 treatment in patients with NSCLC^11^. Whether these associations depend on specific microbial exposures remains unresolved. Our results suggest that host genotype may influence the therapeutic activity of individual bacteria even when their abundance is comparable. Prospective ICB cohorts integrating germline sequencing, longitudinal microbiome profiling, and clinical outcomes would allow direct testing of interactions between NOD2 genotype and microbial composition or function. Such analyses should incorporate functional measurements of microbial immune stimulation, because bacterial abundance or lipoprotein gene content alone may not capture TLR2 activity. Ultimately, defining these host–microbe relationships and elucidating the probiotic mechanisms of action could guide the selection of bacterial strains or microbial products according to the patient’s microbial-sensing capacity, providing a basis for more precise microbiota-based cancer immunotherapy.

**Figure S1.**
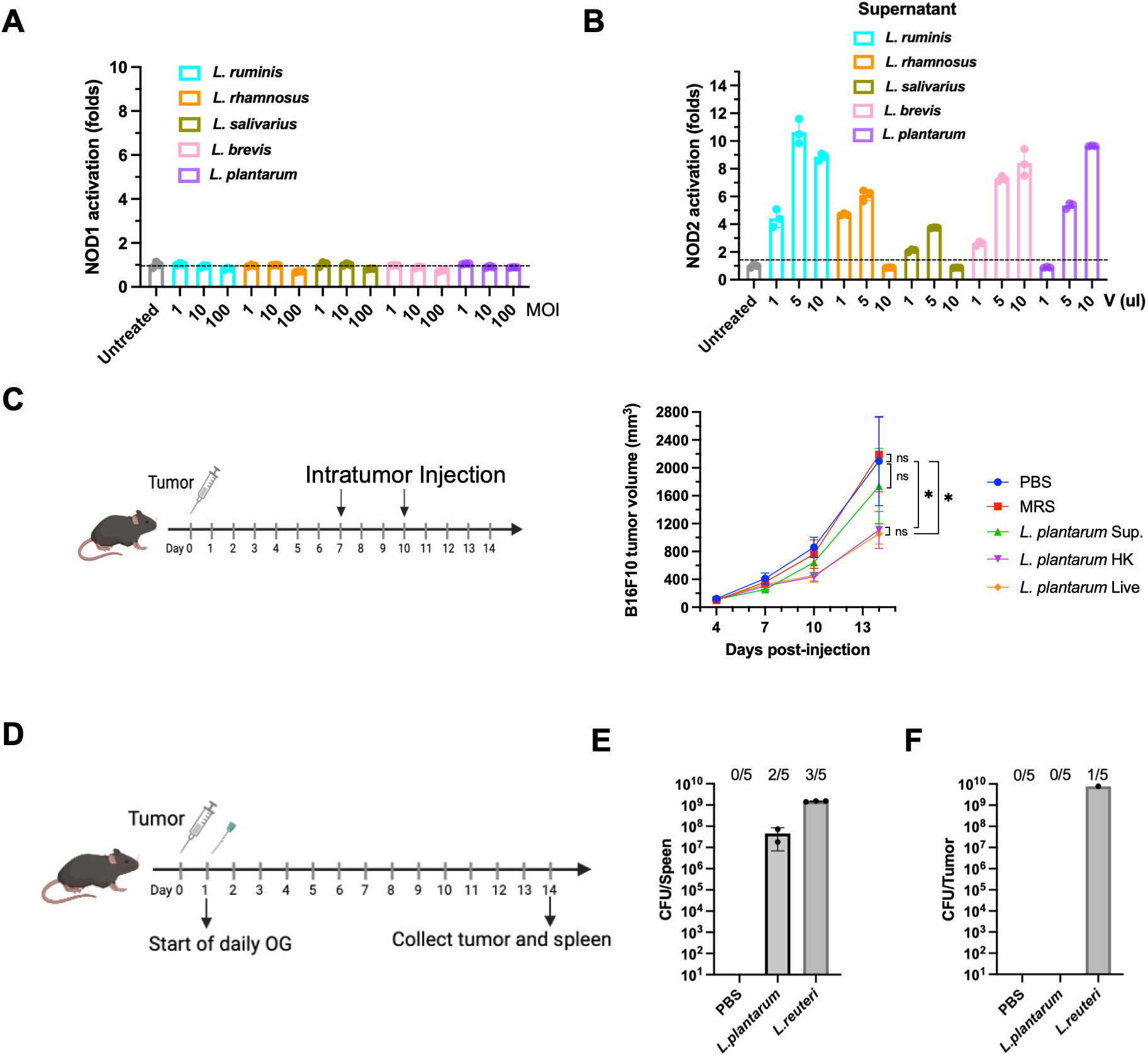
Innate receptor activation by lactobacilli and evaluation of *L. plantarum* antitumor activity and bacterial translocation, related to Figure 1. (A) Activation of NOD2 by supernatants from different *Lactobacillus* species. NOD2-HEKBlue reporter cells were stimulated with the indicated volumes of bacterial culture supernatants in a final volume of 200 μL, and NF-κB activation was quantified. **(B)** Activation of NOD1 by different *Lactobacillus* species. NOD1-HEKBlue reporter cells were stimulated with the indicated *Lactobacillus* species at varying multiplicities of infection (MOIs), corresponding to the indicated bacteria-to-cell ratios, and NF-κB activation was quantified. **(C)** Tumor growth in mice administered intratumorally with sterile-filtered MRS medium, *L. plantarum* culture supernatant, HK *L. plantarum*, or live *L. plantarum* on days 7 and 10 after tumor implantation, as indicated. **(D)** Schematic of the experimental design used to assess bacterial translocation. Mice were orally gavaged with *L. plantarum* daily beginning on day 1. On day 14, tumors and spleens were collected, homogenized, and plated on Lactobacillus Selection (LBS) agar. **(E, F)** Quantification of viable *L. plantarum* recovered from spleens **(E)** and tumor **(F)** by colony-forming unit (CFU) enumeration.

**Figure S2.**
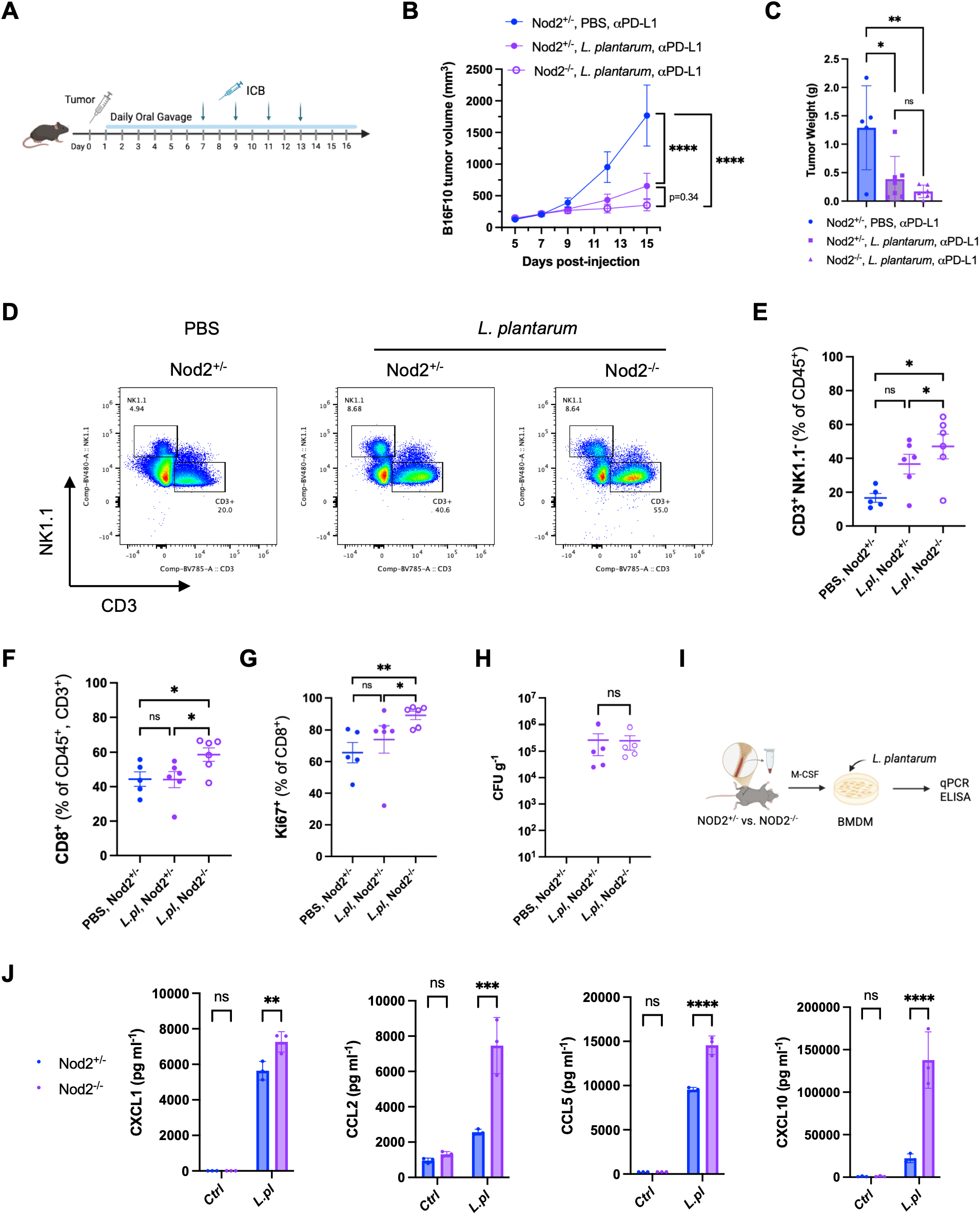
**NOD2 deficiency enhances *L. plantarum*–induced antitumor immunity and myeloid chemokine responses**. **(A–H)** *Nod2^+/-^*and *Nod2^-/-^* littermates bearing B16F10 tumors received daily oral administration of PBS or *L. plantarum*. Anti–PD-1 treatment was initiated on day 7 and administered every other day. **(A)** Schematic of the experimental design. **(B)** B16F10 tumor growth in *Nod2^+/-^* mice treated with PBS and in *Nod2^+/-^*and *Nod2^-/-^* mice treated with *L. plantarum*, all in combination with anti–PD-1. **(C)** Tumor weights measured at the experimental endpoint on day 15. **(D)** Representative flow-cytometry plots showing the identification of CD3^+^NK1.1^−^ T cells and CD3^−^NK1.1^+^ NK cells among live CD45^+^ tumor-infiltrating immune cells. **(E)** Frequency of CD3^+^NK1.1^−^ T cells among live CD45^+^ tumor-infiltrating immune cells. **(F)** Frequency of CD8^+^ T cells among live CD45^+^CD3^+^ tumor-infiltrating T cells. **(G)** Frequency of Ki67^+^ cells among tumor-infiltrating CD8^+^ T cells. **(H)** Fecal abundance of *L. plantarum* in *Nod2^+/-^*and *Nod2^-/-^* mice, quantified as colony-forming units per gram of feces. **(I)** Schematic of the *ex vivo* macrophage-stimulation experiment. Bone marrow cells isolated from *Nod2*^+/−^ and *Nod2*^−/−^ littermates were differentiated with M-CSF for 7 days. The resulting bone marrow-derived macrophages (BMDMs) were stimulated overnight with *L. plantarum*, after which culture supernatants were collected for cytokine and chemokine analysis. **(J)** Concentrations of CXCL1, CCL2, CCL5, and CXCL10 in culture supernatants from unstimulated or *L. plantarum*–stimulated *Nod2^+/-^*and *Nod2^-/-^* BMDMs, measured using a multiplex immunoassay. Tumor-growth data in **(B)** are presented as mean ± SEM and were analyzed using a mixed-effects model followed by Tukey’s multiple-comparisons test. Data in **(C)** and **(E–G)** were analyzed using one-way ANOVA followed by Tukey’s multiple-comparisons test. Data in **(H)** were analyzed using a two-sided unpaired *t* test. Data in **(J)** were analyzed using two-way ANOVA followed by Tukey’s multiple-comparisons test. Each symbol in **(C)** and **(E–H)** represents an individual mouse. For the ex vivo experiments, [n] independent biological replicates were analyzed. Data are presented as mean ± SEM. *\*P < 0.05; **P < 0.01; ***P < 0.001; ****P < 0.0001; ns, not significant*.

**Figure S3.**
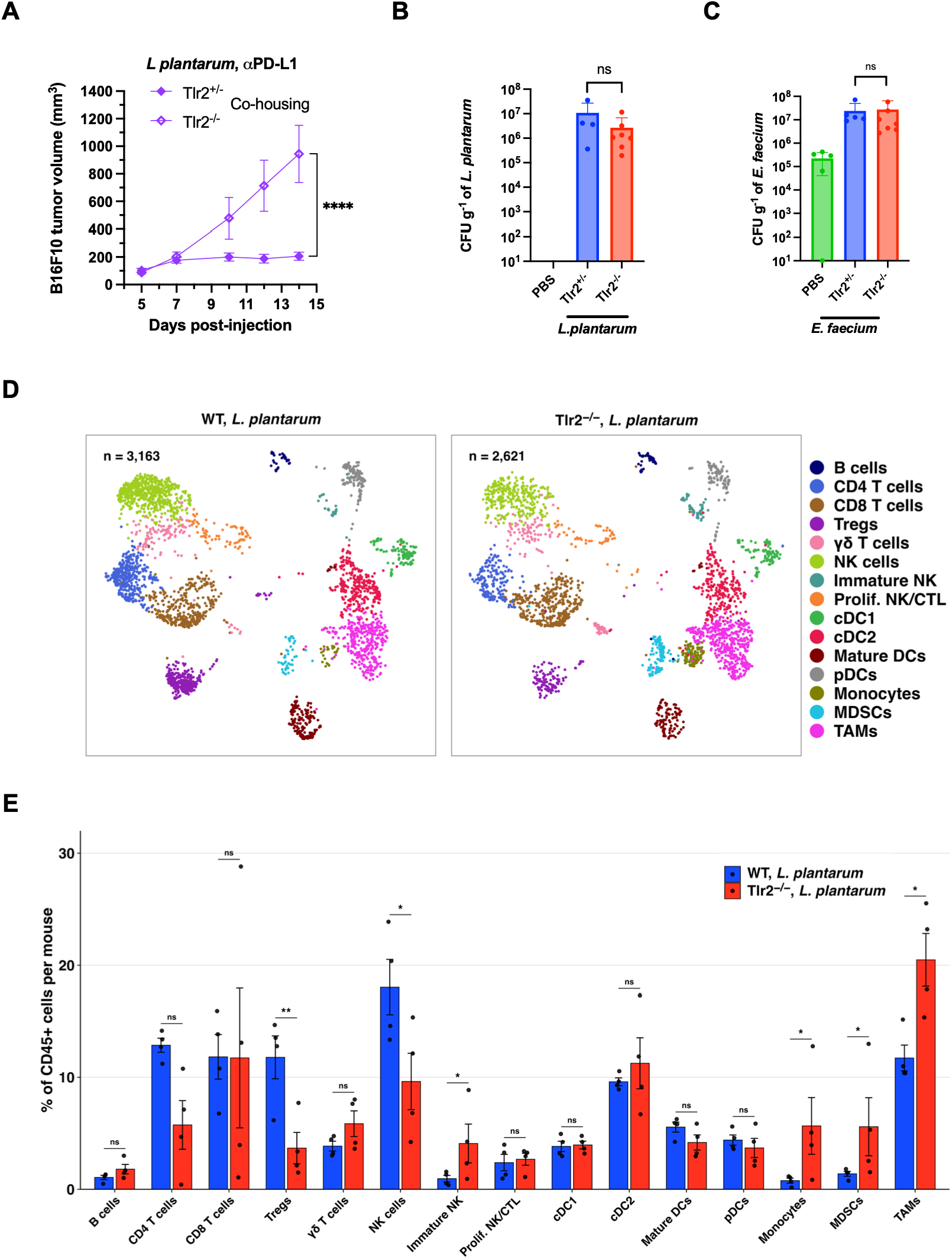
**TLR2 deficiency alters the intratumoral immune landscape without affecting intestinal bacterial abundance**. **(A)** B16F10 tumor growth in cohoused *Tlr2*^+/−^ and *Tlr2*^−/−^ mice receiving *L. plantarum* by oral gavage in combination with anti–PD-L1. Mice were cohoused for 10 days before tumor implantation, and oral administration of *L. plantarum* was initiated on day 1 after tumor implantation. **(B)** Fecal abundance of *L. plantarum* in PBS-treated mice and in *L. plantarum*–treated *Tlr2*^+/−^ and *Tlr2*^−/−^ littermates from the experiment shown in Figure 3A–C. Fecal samples were plated on Lactobacillus-selective agar, and bacterial abundance was quantified as colony-forming units per gram of feces. **(C)** Fecal abundance of *E. faecium* in PBS-treated mice and in *E. faecium*–treated *Tlr2*^+/−^ and *Tlr2*^−/−^ littermates from the experiment shown in Figure 3A and 3D. Fecal samples were plated on HiCrome Enterococci agar, and bacterial abundance was quantified as colony-forming units per gram of feces. **(D)** Uniform manifold approximation and projection (UMAP) of 5,784 tumor-infiltrating CD45+ immune cells from *L. plantarum*–treated WT and *Tlr2*^−/−^ mice, split by genotype and colored by annotated cell type. The analysis included 3,163 cells from WT mice and 2,621 cells from *Tlr2*^−/−^ mice. Both plots use the same integrated embedding and cell-type color scheme. Cell identities were assigned using canonical marker expression. **(E)** Mouse-level proportions of all 15 annotated tumor-infiltrating immune populations, expressed as a percentage of total CD45+ cells in *L. plantarum*–treated WT and *Tlr2*−/− mice. Bars indicate group means, error bars indicate SEM, and points represent individual mice (n = 4 mice per genotype). Raw two-sided *P* values were calculated using propeller with limma-moderated *t* tests applied to logit-transformed per-mouse proportions. Benjamini-Hochberg-adjusted *q* values were calculated across all 15 populations. No comparison reached *q* < 0.05; therefore, raw *P* and adjusted *q* values are displayed directly, and the observed differences should be interpreted as exploratory. Tumor-growth data are presented as mean ± SEM and were analyzed using a mixed-effects model followed by Tukey’s multiple-comparisons test. Bacterial-abundance data were analyzed using one-way ANOVA followed by Tukey’s multiple-comparisons test. Each symbol in **(B)**, **(C)**, and **(E)** represents an individual mouse. *\*P < 0.05; **P < 0.01; ***P < 0.001; ****P < 0.0001; ns, not significant*.

**Figure S4.**
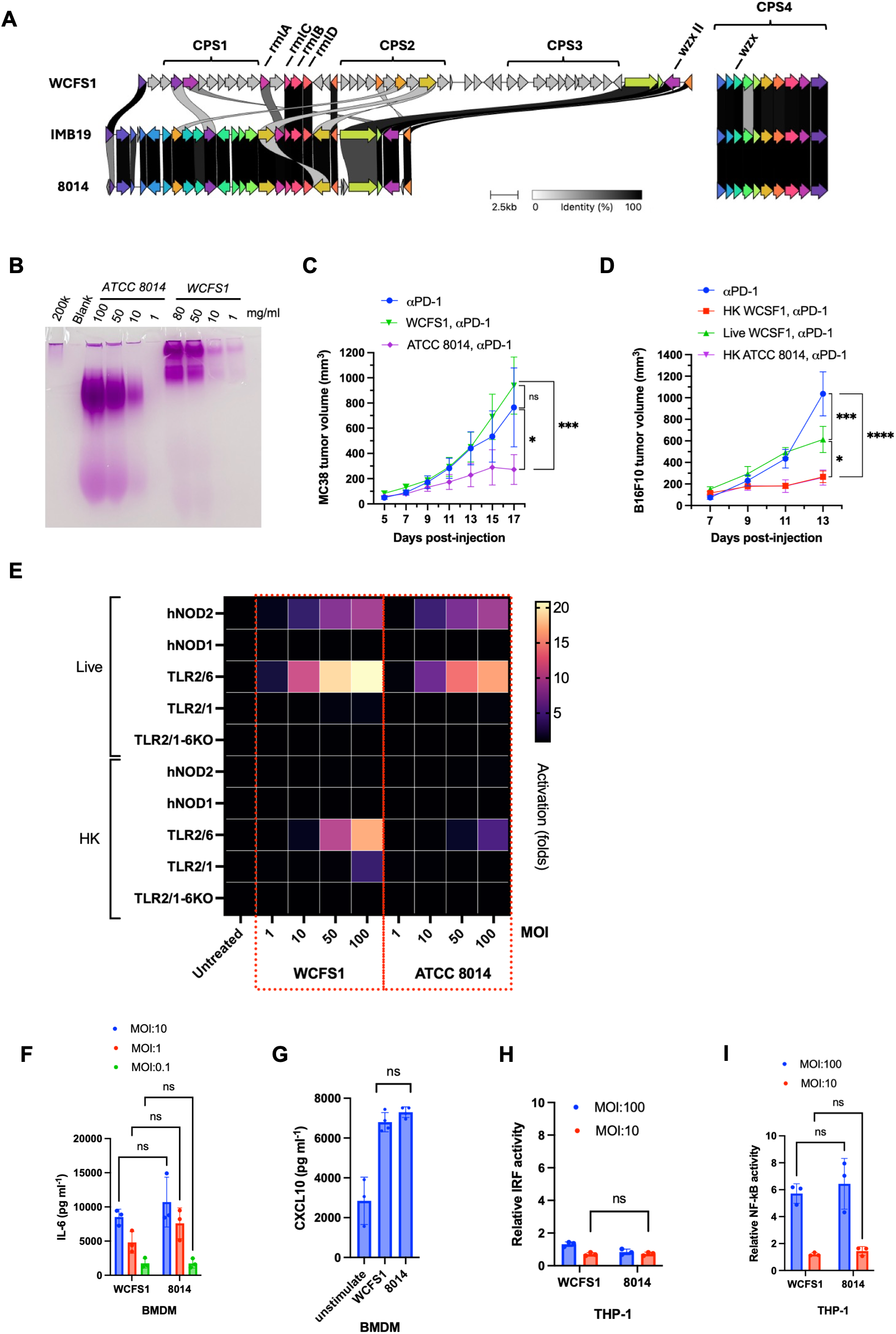
Comparative CPS organization and immune-stimulatory activity of *L. plantarum* strains. **(A)** Comparative organization of capsular polysaccharide (CPS) gene clusters in *L. plantarum* strains WCFS1, IMB19, and 8014. WCFS1 contains four CPS clusters (CPS1–CPS4) and the complete rhamnose-biosynthesis gene set (*rmlA–rmlD*), whereas IMB19 and 8014 retain CPS4 and *rmlA–rmlD*. Gene-cluster alignment was generated using Clinker with a minimum sequence-identity threshold of 30%. Gray shading indicates amino acid sequence identity between homologous genes. **(B)** Gel-based profiling of CPS isolated from *L. plantarum* 8014 and WCFS1 at the indicated concentrations. WCFS1 exhibited higher-molecular-weight CPS species than 8014. **(C)** MC38 tumor growth in mice receiving anti–PD-1 alone or daily oral administration of live *L. plantarum* WCFS1 or 8014 in combination with anti–PD-1. **(D)** B16F10 tumor growth in mice receiving anti–PD-L1 alone or live or heat-killed (HK) *L. plantarum* WCFS1 or 8014 in combination with anti–PD-L1. **(E)** Heatmap summarizing the activation of human NOD1, NOD2, TLR2/6, TLR2/1, and TLR2 KO-TLR1/6 reporter cells by live or heat-killed *L. plantarum* WCFS1 and ATCC 8014 at the indicated MOIs. Color indicates fold activation relative to untreated cells. **(F)** IL-6 concentrations in culture supernatants from mouse BMDM stimulated with *L. plantarum* WCFS1 or 8014 at the indicated MOIs. **(G)** CXCL10 concentrations in culture supernatants from mouse BMDM unstimulated or stimulated with *L. plantarum* WCFS1 or ATCC 8014. **(H)** Relative IRF reporter activity in THP-1 cells stimulated with *L. plantarum* WCFS1 or ATCC 8014 at the indicated MOIs. **(I)** Relative NF-κB reporter activity in THP-1 cells stimulated with *L. plantarum* WCFS1 or 8014 at the indicated MOIs. Tumor-growth data in **(C)** and **(D)** are presented as mean ± SEM and were analyzed using a mixed-effects model followed by Tukey’s multiple-comparisons test. Data in **(F)–(I)** are presented as mean ± SEM; each symbol represents an independent biological replicate. *\*P < 0.05; **P < 0.01; ***P < 0.001; ****P < 0.0001; ns, not significant*.

**Figure S5.**
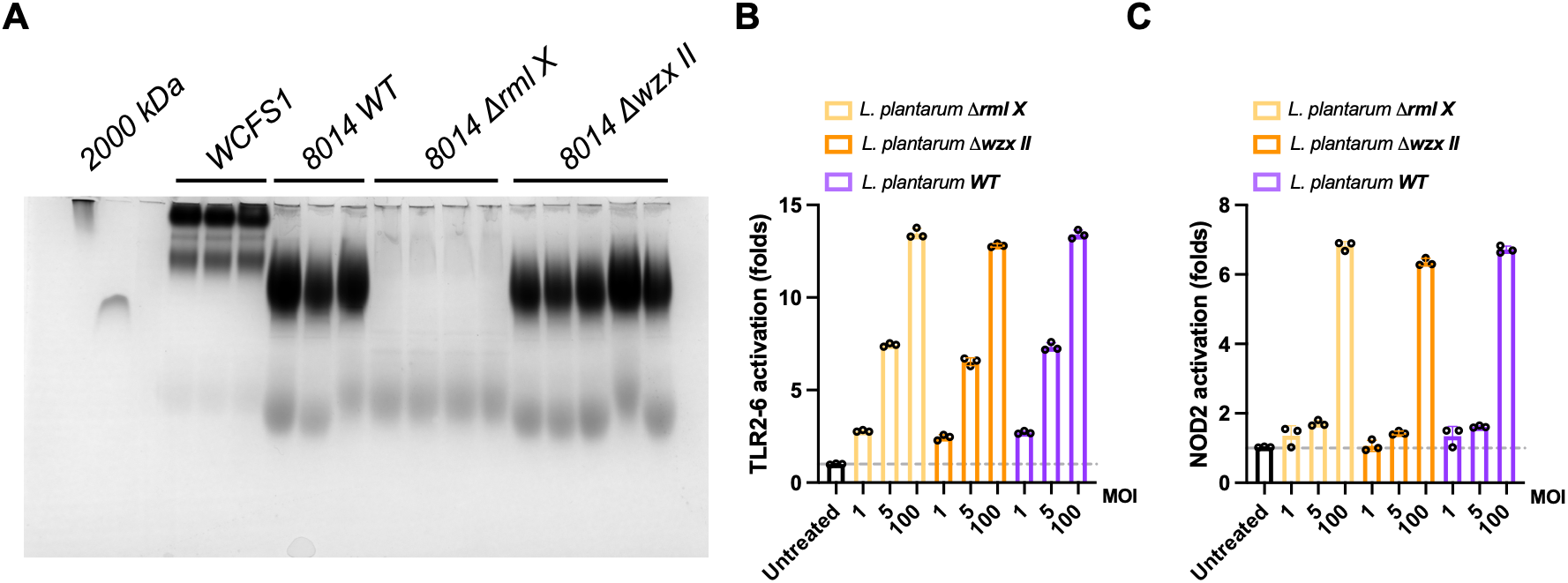
Disruption of CPS biosynthesis does not substantially alter TLR2/6 or NOD2 activation by *L. plantarum*. **(A)** Profiles of surface-associated polysaccharides extracted from *L. plantarum* strains and CPS mutants. Crude polysaccharide extracts (50 mg/ml) were separated on a polyacrylamide gel and stained with Periodic acid-Schiff stain. 2000 kDa Dextran (1 mg/ml) was used as a high molecular weight (HMW) carbohydrate control; Horseradish Peroxidase (1 mg/ml), a heavily glycosylated glycoprotein, was used as a positive control; a non-glycosylated protein, Soybean Trypsin Inhibitor (1 mg/ml), was used as a negative control. **(B)** NF-κB responses of HEK-Blue hTLR2-6 and hNOD2 cells to live *L. plantarum* 8014 and the indicated CPS-biosynthesis mutants. MOI is indicated below. Activation is expressed as fold change relative to vehicle control. Colorimetric QUANTI-Blue detection assay (InvivoGen). Data are mean ± S.D., n = 3.

**Figure S6.**
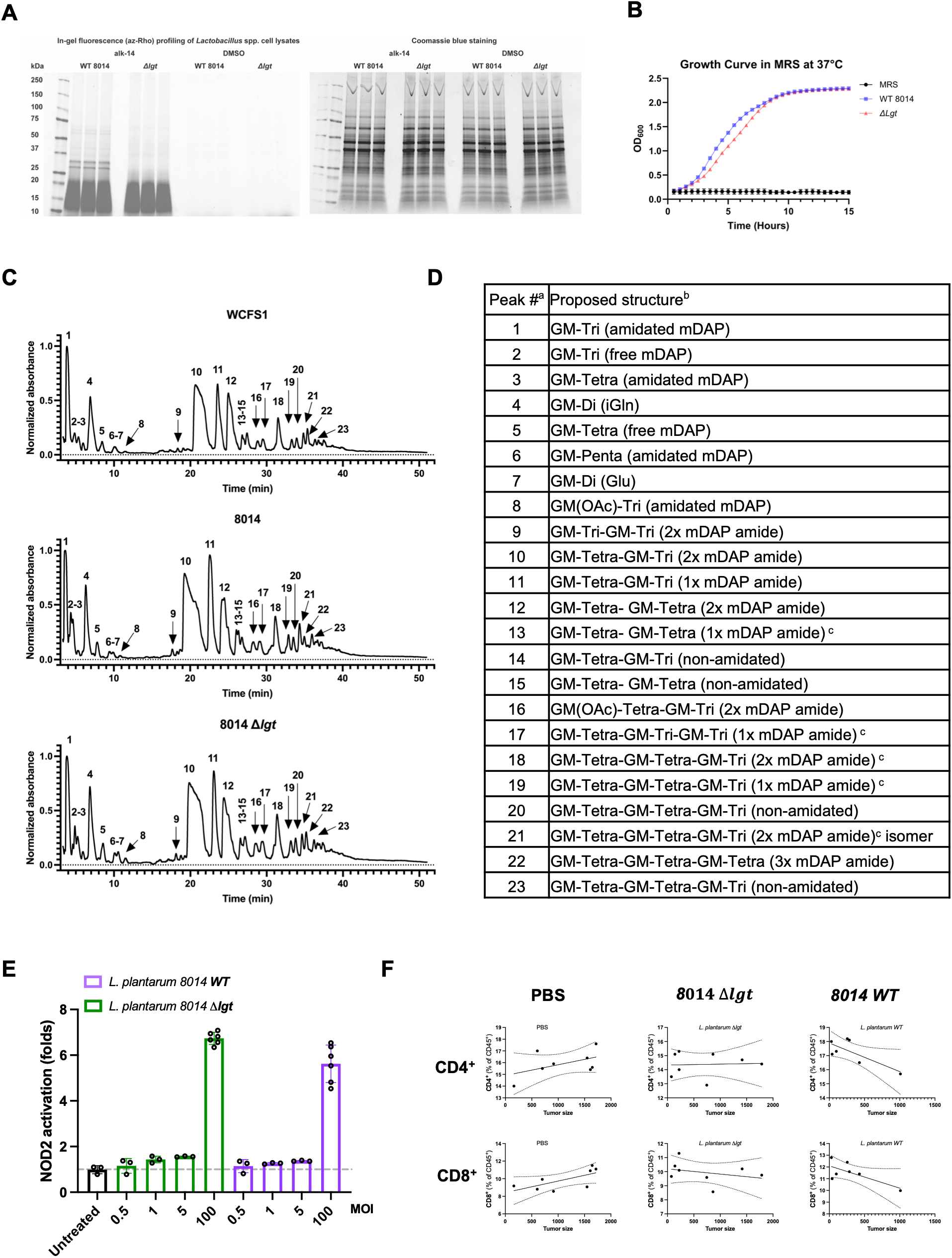
Characterization of the *L. plantarum* Δ*lgt* mutant and its immunostimulatory activity. Related to. Figure 4**. (A)** Lipoprotein profiling of *L. plantarum* ATCC 8014 WT and Δ*lgt* cell lysates. Lipoproteins were metabolically labeled with Alk-14 and detected by in-gel fluorescence following click-chemistry conjugation to azido-rhodamine (az-Rho). Coomassie blue staining shows total protein loading. DMSO-treated samples served as labeling controls. **(B)** Growth of *L. plantarum* ATCC 8014 WT and Δ*lgt* in MRS medium at 37°C, monitored by optical density at 600 nm (OD₆₀₀). **(C)** Representative LC–MS chromatograms of mutanolysin-digested peptidoglycan isolated from sacculi of *L. plantarum* WCFS1, ATCC 8014, and ATCC 8014 Δ*lgt* (top to bottom). **(D)** Composition of peptidoglycan isolated from *L. plantarum* sacculi. ^a^ Peak numbers refer to a. ^b^ GM, disaccharide (GlcNAc-MurNAc); GM-Tri, disaccharide dipeptide (L-Ala-D-iGln or L-Ala-D-Glu); GM-Tri, disaccharide tripeptide (L-Ala-D-iGln-mDAP); GM-Tetra, disaccharide tetrapeptide (L-Ala-D-iGln-mDAP-D-Ala); GM-Penta, disaccharide pentapeptide (L-Ala-D-iGln-mDAP-D-Ala-D-Ala). ^c^ The assignment of the amide and the carboxylate functions to either peptide stem is arbitrary. Masses and retention time of peptidoglycan fragments are in Table S2. **(E)** NOD2 reporter activation by *L. plantarum* ATCC 8014 WT and Δ*lgt* at the indicated MOIs. Reporter activity is shown as fold activation relative to untreated cells. **(F)** Correlations between tumor size and the frequencies of splenic CD4⁺ T cells (top) or CD8⁺ T cells (bottom), expressed as percentages of CD45⁺ cells, in mice treated with PBS, *L. plantarum* Δ*lgt*, or *L. plantarum* WT. Solid lines indicate linear regression fits, and dashed lines indicate 95% confidence intervals. Each symbol represents an individual mouse.

**Figure S7.**
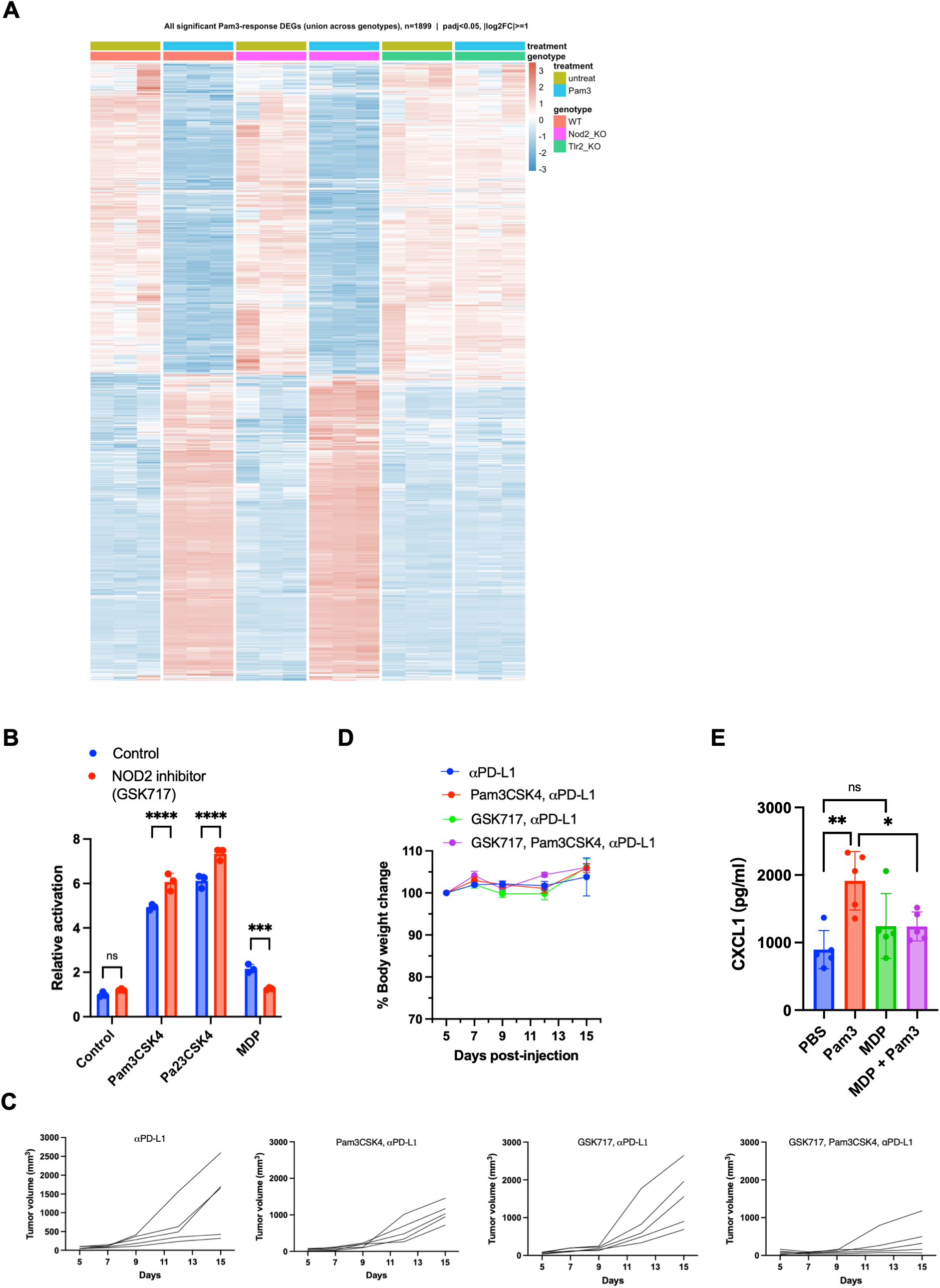
Transcriptional and pharmacological characterization of NOD2 regulation of TLR2 agonist responses. Related to. Figure 5**. (A)** Heatmap showing the union of genes differentially expressed in response to Pam3CSK4 in WT, *Nod2^-/-^*, or *Tlr2^-/-^* BMDMs. Rows represent differentially expressed genes, and columns represent biological replicates from untreated and Pam3CSK4-stimulated BMDMs of each genotype. Expression values are shown as row-scaled normalized expression. **(B)** Relative reporter activity in RAW macrophages pretreated with vehicle or the NOD2 inhibitor GSK717 and subsequently stimulated with Pam3CSK4, Pam2CSK4, or MDP. **(C)** Individual B16F10 tumor-growth curves from the experiment shown in Figure 5F. Each line represents an individual mouse receiving anti–PD-L1 alone or in combination with Pam3CSK4, GSK717, or both compounds. **(D)** Changes in body weight during the B16F10 tumor experiment shown in Figure 5F. Body weight is expressed as a percentage of the weight measured at the beginning of treatment. **(E)** Serum CXCL1 concentrations in mice receiving PBS, Pam3CSK4, MDP, or combined MDP and Pam3CSK4 treatment, as determined by ELISA. Data are presented as mean ± SEM. Each symbol represents an independent biological replicate or an individual mouse. Statistical comparisons in **(B)** were performed using two-way ANOVA followed by Šídák’s multiple-comparisons test, and comparisons in **(E)** were performed using one-way ANOVA followed by Tukey’s multiple-comparisons test. *\*P < 0.05; **P < 0.01; ***P < 0.001; ****P < 0.0001; ns, not significant*.

**Figure S8.**
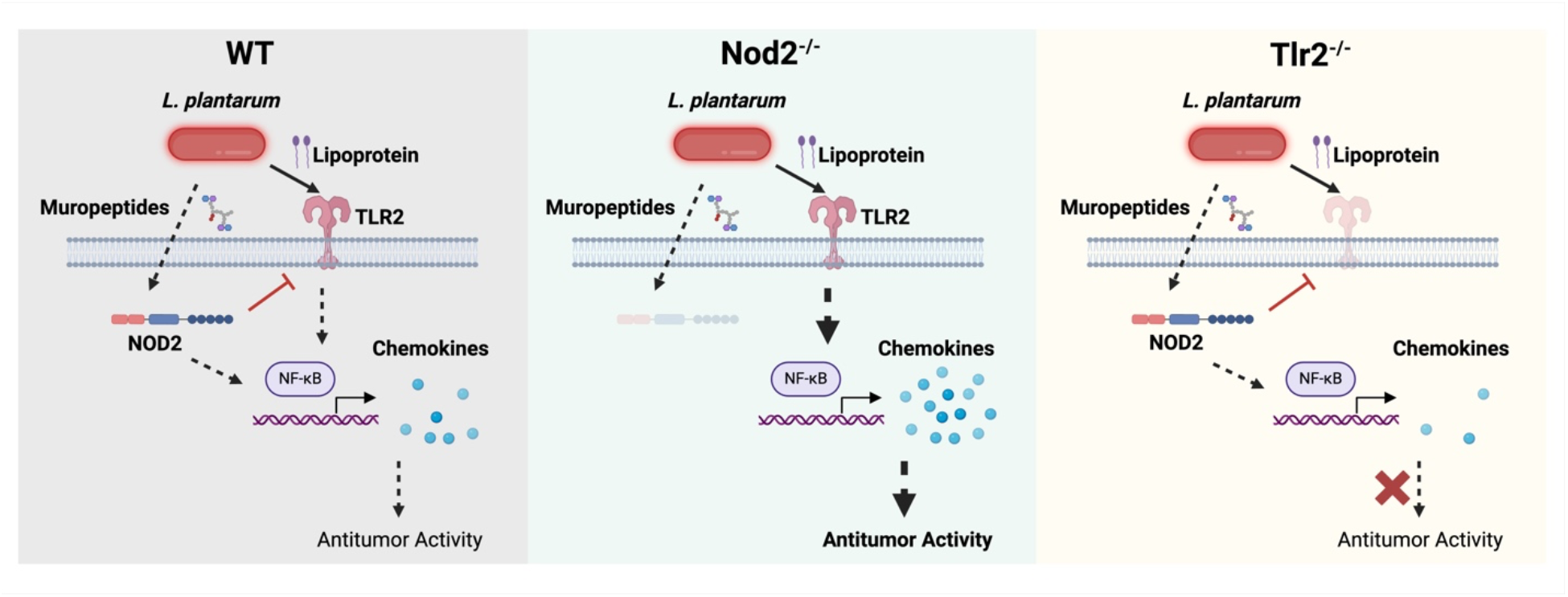
Proposed model of NOD2 regulation of TLR2-dependent, *L. plantarum*–driven antitumor immunity. Cell-associated lipoproteins from *L. plantarum* activate TLR2, inducing chemokine production and immune-stimulatory programs that promote tumor control. *L. plantarum*–derived muropeptides can also activate NOD2. In NOD2- and TLR2-sufficient mice (left), NOD2 limits the magnitude of the TLR2-dependent immune response through an indirect mechanism that remains to be defined. In NOD2-deficient mice (middle), enhanced TLR2-dependent chemokine and immune-stimulatory responses are associated with improved tumor control. In TLR2-deficient mice (right), *L. plantarum* fails to induce an effective antitumor response despite preserved NOD2 sensing, demonstrating that TLR2 is required for *L. plantarum*–mediated tumor control. Dashed regulatory lines denote proposed indirect interactions for which the underlying molecular mechanism has not been established.

## Methods

### Bacterial species

*Lactobacillus rhamnosus* ATCC 21052, *L. ruminis* ATCC 25644, *L. salivarius* ATCC 11741, *L. brevis* ATCC 14869, *L. reuteri* ATCC BAA-2837, and *L. plantarum* WCFS1 and ATCC 8014 were obtained from ATCC. *Enterococcus faecium* Com15 was provided by Michael Gilmore (Harvard Medical School). *Lactobacillus* strains were cultured in de Man, Rogosa and Sharpe (MRS) broth at 37°C under ambient atmosphere, except *L. ruminis* ATCC 25644, which was cultured anaerobically. *E. faecium* was cultured in brain heart infusion (BHI) broth. For colony-forming unit (CFU) enumeration, serial dilutions were plated on MRS, LBS, or HiCrome agar, as appropriate.

### Mice

C57BL/6J mice (stock no. 000664) were obtained from the Scripps Rodent Breeding Colony. *Nod2^-/-^* mice (B6.129S1-*Nod2*^tm1Flv^/J) were obtained from The Jackson Laboratory. *Tlr2^-/-^* mice (B6.129-*Tlr2*^tm1Kir^/J) were provided by Alana Hawkes from the McKay laboratory at Scripps Research. Mice were bred in-house to generate knockout and heterozygous littermates. Mice were housed in autoclaved cages with SaniChip bedding and nesting material on a 12-hour light/dark cycle. Gamma-irradiated chow (LabDiet 5053) and sterile drinking water were provided *ad libitum*. Age- and sex-matched mice were randomly assigned to experimental groups. All animal procedures were approved by the Scripps Research Institutional Animal Care and Use Committee (AUP-21-095) and conducted in accordance with NIH guidelines. For experiments involving bacterial administration, mice were transferred to a containment room before treatment.

### Cell lines and cell culture

HEK-Blue NOD2 (hkb-hnod2), HEK-Blue NOD1 (hkb-nod1), HEK-Blue Null2 (hkb-null2), HEK-Blue TLR2/1 (hkb-htlr21), HEK-Blue TLR2/6 (hkb-htlr26), NF-κB reporter RAW 264.7 (raw-sp), and THP1-Dual (thpd-nfis) cells were obtained from InvivoGen. B16-F10 cells (CRL-6475) were obtained from ATCC, MCA205 cells (SCC173) from MilliporeSigma, and MC38 cells (ENH204-FP) from Kerafast. Cells were cultured at 37°C in 5% CO_2 i_n DMEM or RPMI 1640 supplemented with 10% fetal bovine serum and penicillin–streptomycin. Adherent cells were passaged before reaching 70% confluence, using TrypLE Express where applicable. Cells were used within 10 passages and tested regularly for mycoplasma.

### Reporter activation assays

The assay was followed by manufacturer’s protocol. In brief, reporter cells were seeded in 96-well plate at the cell density 6×10^4^ cells/well. *Lactobacillus* were grown to an OD of ∼0.6 from overnight inoculant, washed with PBS and resuspended in Opti-MEM. Live bacteria were given to HEK-Blue™ cells at indicated multiplicity of infection (MOI) and incubated for 4 hours at 37°C. After 4 hours, HEK-Blue™ cells were carefully washed with PBS and the media was replaced with DMEM supplemented with gentamycin (250 μg/mL) to kill extracellular bacteria. Cells were incubated at 37°C for an additional 16 hours. For secreted metabolites, overnight *Lactobacillus* cultures were spined down at 5000xg for 10 min, and clear supernatants were sterile filtered (0.45 µm). PAMP activation was measured by detecting NF-κB-inducible secreted embryonic alkaline phosphatase expression in the media using colorimetric QUANTI-Blue™ detection assay (InvivoGen). Fold-change of PAMP activation was expressed relative to untreated HEK-Blue™ cells control.

### Generation of NOD2 knock-out THP1-Dual™ cells

The Target Guide Sequence Cloning Protocol provided by the Genome-scale CRISPR Knock-out (GeCKO v2) library (Sanjana NE, Shalem O, Zhang F Nat Methods. 2014 Aug;11(8):783-4. doi: 10.1038/nmeth.3047) was used to construct pLentiCRISPRv2-NOD2, with a NOD2 sgRNA sequence of GAGCACACTCAACCAGCGTG. HEK293T cells were seeded in 100 x 15 mm dishes, and each dish was transfected with 3 µg of envelope vector pMD.2, 6 µg of packaging vector psPAX2, and 6 µg of pLentiCRISPRv2-NOD2 plasmids using Lipofectamine 3000 (Thermo Fisher Scientific). Supernatant was collected 48 hours post-transfection, filtered through a 0.22 µm syringe filter, and mixed with Lenti-X™ viral concentrator (Takara Bio) at a 1:3 Lenti-X™ to supernatant ratio overnight at 4°C. Then, the supernatant was removed, the concentrated viral pellet was resuspended in 1 mL warm RPMI+10% FBS media, and the resuspended viral solution can be used immediately or stored at –80°C. On the day of transduction, 100 µL viral solution was added to 1×10^5^ THP1-Dual™ cells (InvivoGen) in RPMI+10% FBS medium. Medium containing 1.6 µg/mL puromycin was added 48 hours post-transduction, and new media with antibiotics was refreshed every 2 days for 8 days. Lack of activity of NOD2 was assessed by adding high concentrations of MDP and analyzing the resulting NF-κB activation using QUANTI-Blue™ Solution (InvivoGen).

### Generation and stimulation of bone marrow-derived cells

Bone marrow cells were isolated from mice and plated directly in 12-well plates for RNA analysis or 96-well plates for cytokine assays. Cells were cultured in RPMI 1640 containing 10% fetal bovine serum, 1% penicillin–streptomycin, and 20 ng/mL M-CSF to generate bone marrow-derived macrophages (BMDMs). For GM-CSF-derived cultures, 20 ng/mL GM-CSF was used instead of M-CSF. Medium containing the respective growth factor was replaced on day 3. On day 7, the medium was changed to RPMI 1640 containing 10% fetal bovine serum without added growth factor and antibiotics. Cells were stimulated with bacteria or the indicated agonists on day 8.

### RNA extraction and Quantitative RT-PCR

Total RNA was isolated using the RNeasy Plus Kit (Qiagen), and cDNA was synthesized using the iScript Advanced cDNA Synthesis Kit (Bio-Rad). cDNA was diluted 1:10 in nuclease-free water before quantitative PCR with PowerUp SYBR Green Master Mix (Applied Biosystems), according to the manufacturer’s instructions. Expression of *Tlr2*, *Nod2*, *Il6*, *Tnf*, and *Cxcl1* was normalized to *Gapdh* or *Actb*, as indicated, and calculated by the 2^−ΔΔCt^ method. Primer sequences are provided in Table S1.

### Bulk RNA sequencing and analysis

Total RNA was isolated from bone marrow-derived macrophages using the RNeasy Plus Kit (Qiagen). Stranded mRNA libraries were prepared by the Scripps Research Genomics Core using the Trolley RNA-seq workflow, which includes poly(A) selection and random-primed cDNA synthesis. Libraries were sequenced on an Element AVITI instrument with 75-bp paired-end reads, targeting approximately 10 million reads per sample. Sequencing reads were quality controlled, aligned to the mouse reference genome, and summarized as gene-level counts. Differential expression analysis was performed on raw gene counts. Gene set enrichment was performed using the indicated gene collections.

### ELISA and multiplex cytokine analysis

Culture supernatants from BMDMs and mouse serum were collected at the indicated time points. IL-6 (Thermo Fisher Scientific, KMC0061), IL-1β (Thermo Fisher Scientific, 88-7013A-88), TNF-α (Thermo Fisher Scientific, BMS607-3), CXCL1 (R&D Systems, MKC00B-1), and CXCL10 (R&D Systems, DY466-05) were measured by ELISA. BMDM culture supernatants were also analyzed using the Mouse Anti-Virus Response Panel (13-plex) Multi-Analyte Flow Assay (BioLegend, 740621). All assays were performed according to the manufacturers’ instructions.

### Antibiotic pretreatment

For experiments involving antibiotic pretreatment, mice received drinking water containing streptomycin (5 g/L), colistin sulfate (1 g/L), and ampicillin (1 g/L) for one week before bacterial administration, as previously described^15^. Antibiotics were dissolved in autoclaved drinking water and passed through a 0.4-μm filter. Two sterile 50-mL conical tubes fitted with autoclaved no. 6 sipper tubes were provided per cage on sterile Allentown-style wire racks. Antibiotic water was replaced twice during the pretreatment week.

### Tumor inoculation and measurement

Tumor cells were cultured for one week and passaged the day before inoculation. Cells were detached with TrypLE Express (Invitrogen, 12604021), washed, resuspended in cold phosphate-buffered saline (PBS), and counted with a hemocytometer. Mice were anesthetized with 2.5% isoflurane, and the right flank was shaved. Each mouse received a subcutaneous injection of 5 × 10⁵ to 1 × 10⁶ tumor cells in 100 μL PBS into the right flank. Once tumors reached approximately 100 mm³, they were measured every three days with digital calipers. Tumor volume was calculated as length × width² × 0.5, with width defined as the shorter dimension. Mice were euthanized when tumor volume exceeded 2,000 mm³.

### Bacterial administration

Bacteria were administered in drinking water as previously described^15^ or by oral gavage. For oral gavage, *L. plantarum* was cultured overnight without shaking in 10 mL MRS broth, collected, and resuspended in 5 mL sterile phosphate-buffered saline (PBS). Mice received 200 μL of the suspension by oral gavage, corresponding to approximately 10⁹ CFU per mouse.

### *In vivo* treatments

For immune checkpoint blockade experiments, mice received anti–PD-L1 (Bio X Cell, BP0101; 20 μg per mouse) or anti–PD-1 (Bio X Cell, BP0146; 100 μg per mouse) by intraperitoneal (i.p.) injection. Antibodies were diluted in antibody dilution buffer (Bio X Cell) to a final injection volume of 200 μL per mouse. GSK717 was prepared in 10% DMSO, 40% PEG400, and 50% water (v/v/v) and administered by i.p. injection at 200 μg per mouse (approximately 10 mg/kg) in a volume of 200 μL. Pam3CSK4 was dissolved in water and administered by i.p. injection at 30 μg per mouse. Control mice received the corresponding vehicle.

### Fecal bacterial burden and bacterial translocation

Fecal samples were collected aseptically, weighed, resuspended in sterile PBS, homogenized with sterile pestles, and serially diluted in PBS. Dilutions were plated as drops on selective agar. Tumors and spleens were collected aseptically, weighed, and homogenized by passage through a 100-μm cell strainer. Samples were incubated anaerobically at 37°C for 3 h, followed by the addition of 5 mL MRS broth and overnight incubation. Enriched samples were serially diluted and plated on MRS or LBS agar. Plates were incubated at 37°C under ambient atmosphere for 24–48 h, and colonies were counted manually.

### Phylogenetic analysis

To identify the population of *L. plantarum* we downloaded all complete genomes on NCBI (n= 1100 as of June 2025). All assemblies were annotated using the run_prokka function in Panaroo (v1.2.10) with clean-mode set to strict, which annotates each sample with the same gene model using Prokka (v1.14.6). The pangenome was defined using Panaroo (v1.2.10), which utilizes a pangenome graph-based approach for clustering. Core genes were defined as genes present in >99% of strains, with accessory in at least >1%. From the pangenome, functional annotation was assigned using eggNOG-mapper (v2.1.2) with default Diamond mode. A maximum-likelihood phylogenetic tree using the alignment of core genes (core_gene_alignment_filtered.aln) was inferred using IQ-TREE (v2.1.4) with a general time-reversible (GTR+G4) substitution model and 1,000 bootstrap replicates. All figures were generated in R (v.4.3.0, https://www.r-project.org/) using tidyverse (v.1.3.1), patchwork (v.1.1.1), ggtree (v.3.8.2), and ggnewscale (v.0.4.5).

Generation of *L. plantarum* ATCC 8014 mutants.

Main challenges of lactobacilli gene editing are difficulties in transforming them with exogenous DNA, due to their thick and complex cell wall structures, presence of native restriction-modification systems, and lack of strain specific approaches. Majority of gene editing has been done in *L. plantarum* WCFS1 strain. Here we show an improved gene editing approach for *L. plantarum* 8014. Targeted mutants were generated by adapting published RecET-mediated recombineering methods^48–50^. The chloramphenicol resistance marker of pLH01 (Addgene #117261), which encodes the phage recombinases RecE and RecT, was replaced with an erythromycin resistance marker. The modified plasmid was propagated in dam⁻/dcm⁻ *E. coli* and electroporated into *L. plantarum* ATCC 8014. For each target locus, a linear recombination cassette containing the chloramphenicol acetyltransferase gene (*cat*) flanked by approximately 1-kb homology arms was assembled by overlap PCR. Cells carrying the modified pLH01 plasmid were grown in MRS medium at 30°C. When the culture reached an OD₆₀₀ of 0.6, SppIP (0.5 μg/mL; MAGNSSNFIHKIKQIFTHR; GenScript) was added to induce recombinase expression. Freshly prepared electrocompetent cells were washed with 10 mM MgCl₂ and 10% glycerol containing 0.5 M sucrose. Approximately 1 μg of purified recombination cassette was electroporated into 50–60 μL of cells in a 1-mm cuvette using a Bio-Rad Gene Pulser II (1.3 kV, 25 μF, 200 Ω). Cells were recovered for 3–4 h at 30°C in MRS medium supplemented with 0.5 M sucrose and 0.1 M MgCl₂, then plated on MRS agar containing erythromycin and chloramphenicol (10 μg/mL each). Mutants were screened by colony PCR and verified by whole-genome sequencing.

### Bacterial lipoprotein profiling

Metabolic labeling of *L. plantarum* lipoproteins was adapted from a published protocol^28^. Overnight cultures were diluted 1:50 in fresh MRS medium and grown statically to an OD₆₀₀ of approximately 0.4. Cultures were then incubated for 1 h at 30°C with 20 μM alkynyl palmitic acid (alk-14; Cat. # A491321) or an equivalent volume of DMSO. Bacteria were collected by centrifugation, washed twice with cold DPBS, and lysed by bead beating in PBS containing an EDTA-free protease inhibitor cocktail. Lysates were clarified by centrifugation, and protein concentrations were determined by BCA assay. For copper-catalyzed azide–alkyne cycloaddition (CuAAC), 50–100 μg of protein in 90 μL was mixed with 10 μL of 10× reagent mixture to give final concentrations of 0.1 mM azide-rhodamine, 1 mM TCEP, 0.1 mM TBTA, and 1 mM CuSO₄.

Reactions were incubated for 1 h at room temperature. Proteins were precipitated with four volumes of methanol, washed twice with methanol, air-dried, and resuspended in SDS sample buffer. Samples were heated at 95°C for 10 min and separated on 4%–20% SDS-PAGE gels. Rhodamine-labeled proteins were visualized by fluorescence imaging, followed by Coomassie staining to assess total protein loading.

### Profiling of surface-associated polysaccharides

Surface-associated polysaccharides were extracted from *L. plantarum* ATCC 8014 using a published protocol with minor modifications^51^. Overnight cultures were diluted 1:50 in MRS medium and grown statically at 37°C to an OD₆₀₀ of 0.8–1.0. Cells were washed twice with PBS and resuspended in buffer containing 25% sucrose, 10 mM Tris, and 0.05% sodium azide. Lysozyme (1 mg/mL) and mutanolysin (10 U/mL) were added, and samples were incubated for 16 h at 37°C with gentle rocking. Following centrifugation, supernatants were treated with RNase (100 μg/mL), DNase (10 U/mL), MgCl₂ (2.5 mM), and CaCl₂ (0.5 mM) for 4 h at 37°C, followed by pronase (50 μg/mL) for 16 h at 37°C. Samples were extracted with chloroform, and the aqueous phase was collected. Polysaccharides were precipitated with ethanol overnight at −80°C, air-dried, and resuspended in water at 50 mg/mL. Aliquots (12 μL) were separated on 4%–20% polyacrylamide gels at 120 V for 1 h and stained with periodic acid–Schiff reagent (Pierce Glycoprotein Staining Kit, Thermo Fisher Scientific). Dextran (2,000 kDa), horseradish peroxidase, and soybean trypsin inhibitor were included as carbohydrate, glycoprotein, and nonglycosylated protein controls, respectively.

### Peptidoglycan isolation and LC–MS analysis

Peptidoglycan was isolated and analyzed as previously described with minor modifications^16,52,53^. Overnight bacterial cultures were subcultured in MRS medium and grown to an OD₆₀₀ of approximately 0.8–1.0. Cell pellets were suspended in 0.25% SDS in 0.1 M Tris-HCl (pH 6.8) and boiled for 20 min. Insoluble cell walls were collected by centrifugation, washed with water to remove SDS, sonicated for 30 min, and treated sequentially with benzonase and trypsin. The material was then treated with 1 M HCl for 4 h at 37°C with shaking, collected by centrifugation, and washed with water until the wash reached pH 5–6. Purified peptidoglycan was digested with mutanolysin from *Streptomyces globisporus* (10 KU/mL; Sigma) in 10 mM sodium phosphate buffer (pH 4.9) for 16 h at 37°C. After heat inactivation and centrifugation, soluble muropeptides in the supernatant were reduced with sodium borohydride in 0.25 M boric acid (pH 9) for 1 h at room temperature. The reaction was quenched with orthophosphoric acid and adjusted to pH 2–3. Muropeptides were analyzed on an Agilent 1290 Infinity II LC/MSD system equipped with a Poroshell 120 EC-C18 column (3 × 150 mm, 2.7 μm). Samples were eluted at 0.5 mL/min with water containing 0.1% formic acid (A) and acetonitrile containing 0.1% formic acid (B), using 2% B for 0–5 min followed by a linear increase to 10% B over 5–65 min. Elution was monitored at 205 nm, and masses were acquired in API-ES scan mode (*m/z* 200–2,500). The relative abundance of each assigned muropeptide was calculated as its integrated chromatographic peak area divided by the sum of the areas of all assigned peaks (Table S2).

### Cell isolation and flow cytometry

Tumors, spleens, and tumor-draining lymph nodes (tdLNs) were collected at the indicated endpoints. Tumors were trimmed of excess stromal tissue and dissociated in RPMI containing Liberase TM (1.5 U/mL; Roche, 5401119001), DNase I (0.2 mg/mL; Worthington, LS002006), and 6.35-mm ceramic spheres (MP Biomedicals, 116540424-CF). Samples were shaken gently at 37°C for 30 min and passed through a 70-μm strainer. Spleens and tdLNs were mechanically dissociated through strainers using a syringe plunger. Tumor and spleen suspensions were treated with red blood cell lysis buffer (Thermo Fisher Scientific, 00-4333-57) for 1–2 min at room temperature. Cell suspensions were then passed through 100-μm nylon mesh and transferred to 96-well plates.

Cells were stained with Zombie Yellow viability dye (1:1,000; BioLegend, 423103) for 20 min at room temperature in the dark. After washing, Fc receptors were blocked with TruStain FcX anti-mouse CD16/32 (0.5 μL per sample; BioLegend, 101319) for 30 min at room temperature. Cells were stained on ice for 30 min with antibodies against CD4 (BUV496, GK1.5, BD Biosciences, 612953), CD44 (PerCP-Cy5.5, IM7, BioLegend, 103031), Sca-1 (BUV395, D7, BD Biosciences, 563990), CD69 (PE, H1.2F3, BioLegend, 104507), and PD-1 (BV711, CD279, BD Biosciences, 135231). Cells were washed, fixed, and permeabilized overnight at 4°C using the FoxP3/Transcription Factor Staining Buffer Set (Thermo Fisher Scientific, 00-5523-00). After three washes with permeabilization buffer, cells were incubated with 10% rat serum (Thermo Fisher Scientific, 24-5555-94) for 20 min on ice and stained for 20 min on ice with antibodies against CD45 (APC-Fire 750, 30-F11, BioLegend, 103153), CD3 (BV785, 17A2, BioLegend, 100232), NK1.1 (BV480, PK136, BD Biosciences, 746265), CD8 (PE-Cy7, 53-6.7, BioLegend, 100721), FoxP3 (AF532, FJK-16s, Thermo Fisher Scientific, 58-5773-80), granzyme B (PE-CF594, GB11, BD Biosciences, 562462), and Ki-67 (FITC, SolA15, Thermo Fisher Scientific, 11-5698-82). Data were acquired on a Cytek Aurora spectral flow cytometer and analyzed with FlowJo v10.9.0.

### scRNA-seq sample preparation

B16F10-bearing wild-type, *Nod2^-/-^*, and Tlr*2^-/-^*mice were treated as described for the corresponding *in vivo* experiments. Wild-type and *Nod2^-/-^* mice received oral gavage with PBS or *L. plantarum*; *Tlr2*−/− mice received *L. plantarum* only. Tumors were collected on day 10 after inoculation and processed into single-cell suspensions as described for flow cytometry. Briefly, tumors were enzymatically digested, passed through a 70-μm filter, treated with ACK red blood cell lysis buffer, and washed in cold PBS containing 2% FBS. Cells were stained with anti-CD45 antibody (APC-Fire 750, clone 30-F11; BioLegend, 103153), TotalSeq-C anti-mouse hashtag antibodies (BioLegend), and SYTOX Blue viability dye. Live CD45⁺ cells were sorted on a Bigfoot Spectral Cell Sorter (Thermo Fisher Scientific) using a 100-μm nozzle at 25 psi. Approximately 10,000 live CD45⁺ cells were sorted per sample. Hashtagged cells from individual mice were pooled, and approximately 50,000 cells were targeted for encapsulation on the Chromium Single Cell platform (10x Genomics). Gene expression and feature barcode libraries were prepared using the Chromium GEM-X Single Cell 5′ Reagent Kits v3 and 5′ Feature Barcode Kit v3 (10x Genomics; protocol CG000734).

### scRNA-seq data processing and analysis

Sequencing reads were processed using Cell Ranger (v9.0.1), and gene expression matrices were analyzed in R using Seurat. Cells were assigned to individual mice from hashtag oligonucleotide counts using *HTODemux*. Cells with ambiguous assignments or classified as doublets were excluded. Low-quality cells were removed based on detected gene and transcript counts and the proportion of mitochondrial transcripts. Data were normalized, and highly variable genes were used for principal component analysis and shared nearest-neighbor clustering. Cell types were annotated using established lineage markers; myeloid and lymphoid populations were reclustered to refine cell-type assignments. Differential expression and pathway analyses were performed within the indicated cell types to compare genotype and *L. plantarum* treatment groups.

### Statistical analysis

Statistical analyses were performed using GraphPad Prism 8.0. Two-group comparisons were analyzed using two-sided unpaired *t* tests or Mann–Whitney tests, as appropriate. Comparisons involving multiple groups were analyzed by one-way or two-way ANOVA followed by the multiple-comparison test specified in the figure legend. Adjusted *P* values are reported for multiple comparisons. The test used for each experiment is indicated in the corresponding figure legend. *P* < 0.05 was considered statistically significant.

## Acknowledgments

We thank the Scripps Research Department of Animal Resources (DAR), Flow Cytometry, and Genomics Core for technical support.

## Author contributions

X.Z. and H.C.H. conceived the study and developed its conceptual framework. X.Z. performed all *in vivo* experiments, maintained and bred the knockout mouse colonies, conducted all flow cytometry experiments, performed scRNA-seq experiments and most BMDM experiments, and performed the RAW-Blue cell assays and THP-1 assays. N.D. generated the bacterial mutants, performed lipoprotein and CPS profiling, and contributed to reporter and ELISA assays. K.T.F. introduced the bacterial strains and performed reporter assays and peptidoglycan profiling. S.D. established the BMDM assay and contributed to multiplex assays, scRNA-seq experiments and analysis, mouse tissue collection, and flow cytometry troubleshooting. A.M.T. performed bacterial genomic analysis. A.V.-N. assisted with mouse tissue collection. B.A.N. generated the THP-1 NOD2 knockout cells. T.D. performed CPS profiling. X.Z. wrote the original draft. H.C.H. provided scientific input and contributed to manuscript revision, and all authors reviewed and edited the manuscript.

